# Primate Gaze Behavior Reflects Change of Mind in a Naturalistic Navigation Task

**DOI:** 10.64898/2026.10.10.756925

**Authors:** Woochul Choi, Panos Alefantis, Georgios Andreadakis, Dora E. Angelaki

**Affiliations:** Center for Neural Science, New York University, New York, NY, USA; Department of Brain Convergence, Korea Brain Research Institute, Daegu, Republic of Korea; Tandon School of Engineering, New York University, New York, NY, USA

**Author notes:** Correspondence: Woochul Choi and Dora E. Angelaki.

## Abstract

Revising a decision while acting on it is a fundamental aspect of natural behavior, yet the underlying computations are difficult to track in real time. To explore how such belief updates unfold, we tested whether eye movements reveal a subject’s change of mind during a naturalistic navigation task with a binary target choice. Humans and monkeys were required to navigate to one of two goal locations presented either simultaneously or sequentially in a virtual environment where explicit position cues were absent. Our findings reveal that gaze continuously tracks the latent variable, including changes of mind. We found that the target tracking accuracy before the second target is even presented during sequential target trials predicts whether the subject’s navigational goal persists (first target choice) or changes (second target choice). These results underscore the importance of gaze in action planning and changes of mind, supporting the hypothesis that eye movements embody not only the current latent variable but also its uncertainty.

**Research highlights:**

- Humans and monkeys virtually navigated to one of two sequentially presented goals.
- Humans and monkeys prefer a cost-efficient goal, but the introduction of a new goal may sway their initial decisions.
- Eye movements predict change of decision by tracking the chosen goal location.
- Eye movements convey latent information for future navigation planning.

**Results:** 1. Primates virtually navigate and select one of two sequentially presented goals.

2. Sequential goals prompt humans and monkeys to alter their initial decision, reflecting a tradeoff between memory uncertainty and action cost.

3. Eye movements predict change of decision by tracking the invisible target location.

4. Eye movements convey task-related latent variables for navigation.

## Introduction

Our brains make decisions swiftly and adaptively under uncertain and changing conditions. Although seemingly effortless, the underlying computations can be demanding especially when the latent states are continuous, time-varying and not directly observable. To investigate flexible decision-making, a large body of studies examined the neural and behavioral basis of sensory based decisions ^1–7^. However, these paradigms typically impose strong constraints on natural behavior, most notably requiring subjects to maintain central fixation. This restriction stands in sharp contrast to real-world behavior, where observers actively move their eyes to gather information, reduce uncertainty, and support ongoing decisions^8–10^. By limiting exploratory gaze, traditional tasks may fail to engage the full spectrum of embodied mechanisms that underlie natural cognition ^11,12^. To better understand decision-making in naturalistic contexts, it is essential to study behavior under conditions that allow free eye movements during goal-directed tasks—where the brain must actively acquire, update, and act upon uncertain information.

One domain where free eye movement may play a critical role is path integration—where animals navigate to latent goal locations using continuously updated sensory input and self-motion signals ^13–20^. Unlike fixed-fixation tasks, these behaviors require the brain to form and update internal estimates of variables such as position, heading, and goal location, often guided by active visual exploration. Recent studies have employed closed-loop virtual navigation paradigms in which subjects steer toward a briefly flashed target using sparse optic flow cues in virtual reality ^21–25^. These paradigms revealed how primates use continuously evolving sensory information to guide navigation—while freely moving their eyes.

Eye movements in these tasks are not merely epiphenomenal—they provide a continuous window reflecting the internal goal representations. Lakshminarasimhan et al. (2020) showed that primate gaze continuously tracks the remembered goal location even when the target is no longer visible. In human experiments, Stavropoulos et al. (2025) demonstrated that participants continue to align their gaze with the goal location even in the absence of optic flow. These findings suggest that eye movements are essential for building internal beliefs. However, whether gaze also captures the dynamic process of belief updating during decision revision remains unknown. To address this, it is essential to examine gaze behavior in tasks where decisions can evolve within trial, rather than being pre-selected before the action.

Here, we extend this paradigm by introducing a second transient target during ongoing navigation, allowing subjects to revise their initial decisions in response to new sensory evidence. This enables us to investigate how internal beliefs evolve within a single trial, as subjects may change their mind during navigation. Specifically, we test whether eye movements reflect these evolving decision states—beyond simply indicating the chosen goal—by revealing the strength or uncertainty of the underlying belief. If gaze encodes belief uncertainty, then the precision with which subjects track the initial target before the second appears should predict whether they maintain their original decision or switch to the newly introduced alternative.

## Results

### Monkeys and Humans Virtually Navigate to one of two Goals by Integrating Optic Flow

Monkeys and humans performed a closed-loop navigation task in which they used a joystick to steer toward briefly presented targets in a three-dimensional virtual environment. Two circular targets were shown sequentially or simultaneously at pseudo-random positions on a ground plane populated with transiently blinking visual elements, generating optic flow in the absence of stable landmarks (**Figure 1A, B**, see Methods for details ^21,23–25^). After target presentation, subjects navigated based on memory of the locations, using two degrees of freedom—forward and angular velocity—to steer toward one of the targets of their choosing. A reward was delivered upon stopping near the (invisible) target location, with graded feedback based on the final distance from the target center (**Figure 1C**). Eye movements and movement trajectories were recorded continuously throughout the trial.

**Figure 1.**
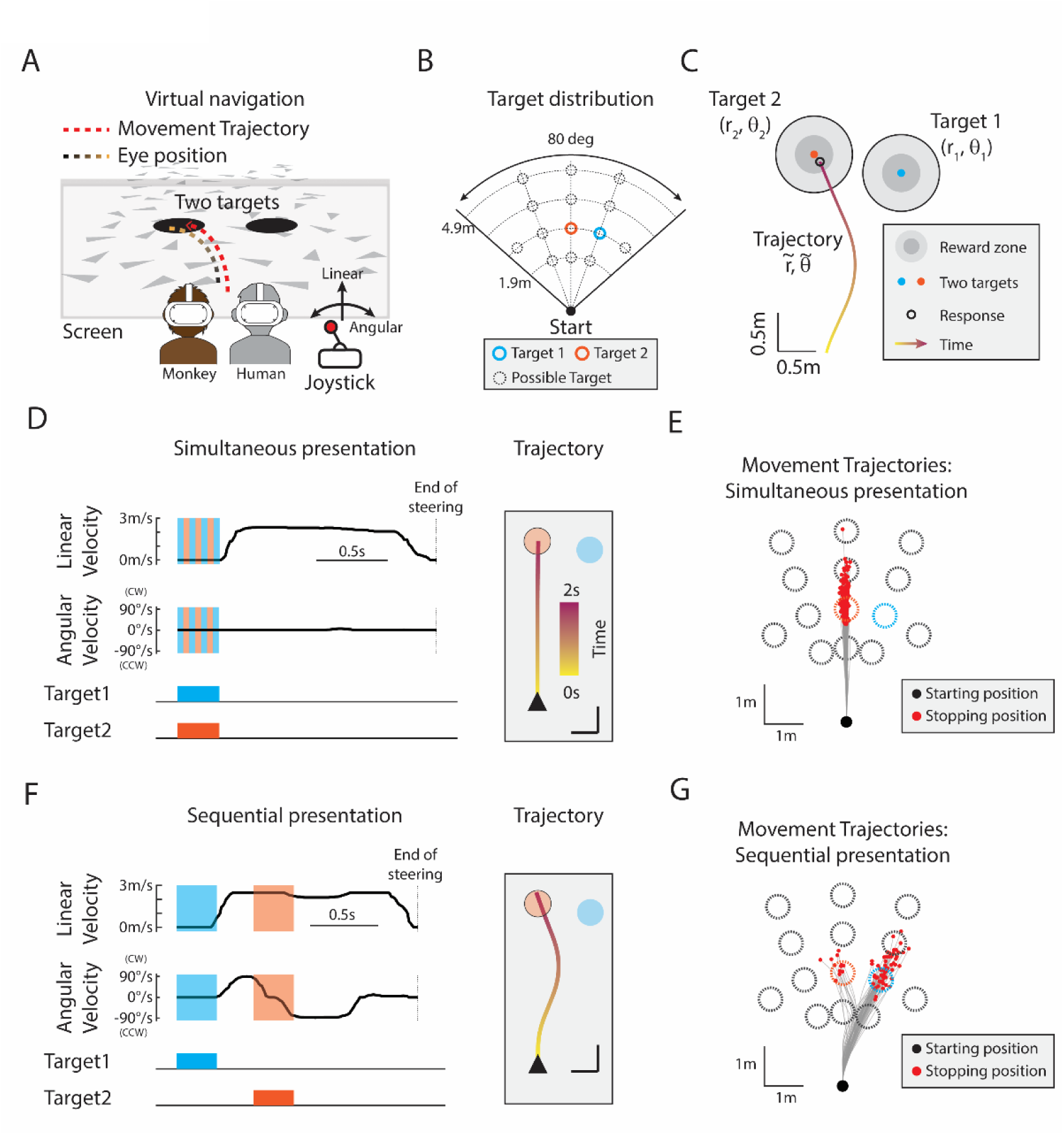
Sequential vs. Simultaneous target presentation tasks. **(A)** Task schematic. Monkeys and humans navigated in a virtual environment using a joystick with two degrees of freedom, corresponding to linear and angular velocities. Subjects steered toward one of two briefly visible targets (black disks) using optic flow cues. **(B)** Overhead view showing one target pair configuration (cyan: first target; orange: second target). See **Supplementary Figure S1** for all target configurations. **(C)** Sample response and reward structure. Subjects received a high reward for landing within 0.25 m of a target center, a smaller reward for landing within 0.25–0.5 m, and no reward otherwise. A representative trajectory is shown alongside the graded reward zones. **(D)** Simultaneous condition: time course of linear and angular velocities. Cyan and orange shading indicate the visibility period for each target. The corresponding movement trajectory is shown in an overhead view (right; scale bar = 0.5 m), using the configuration from panel B. **(E)** Simultaneous condition: representative trajectories and stopping positions. Initial positions are marked with black dots; final stopping points with red dots. Cyan and orange circles indicate the two target locations (same as panel B). **(F)** Sequential condition: time course of linear and angular velocities, formatted as in **(D)**. The corresponding movement trajectory is shown in an overhead view (right; scale bar = 0.5 m), using the configuration from panel B. **(G)** Sequential condition: representative trajectories and stopping positions, formatted as in **(E)**.

Two experimental conditions were interleaved: simultaneous and sequential binary target presentations (**Figure 1D,E** for simultaneous; **Figure 1F,G** for sequential; see **Supplementary Figure S1**). In simultaneous trials, both targets appeared at trial onset and disappeared after 300 ms (**Figure 1D**). Subjects typically committed to one of the two options immediately and maintained their decision throughout navigation without deviating to the alternative (**Figure 1E**). In contrast, sequential trials introduced the first target initially, followed by the second target after a variable delay (Figure 1F). In this condition, subjects frequently adjusted their path during navigation in response to the newly appearing target, reflecting a flexible change of mind (**Figure 1G**). In these trials, the first target appeared at one of three fixed angular positions—straight ahead (0°) or at ±20° eccentricity—at a constant radial distance of 2.9 meters. The second target was then presented at one of five configurations defined relative to the first target: either at three different radial distances along the same angle, or at two different angular positions at a fixed distance, making its location unpredictable (see **Supplementary Figure S1** for details). On average, both humans and monkeys performed well in both conditions: simultaneous (humans: 69% ± 15%; monkey J: 57%; monkey V: 60%) and sequential (humans: 70% ± 15%; monkey J: 56%; monkey V: 54%). Performance in both multi-target conditions exceeded that of single-target controls, with no significant difference between the simultaneous and sequential conditions (see **Supplementary Figure S1**).

### Sequential Target Presentation Biases Navigational Decisions and Reveals Changes of Mind

To examine how subjects made decisions between two competing goals, we analyzed target choice probabilities in simultaneous and sequential presentation trials. Trials were grouped by spatial configuration: either differing in distance at the same angle (**Figure 2A-C**) or differing in angle at the same distance (**Figure 2D-F**). The simultaneous condition provided a baseline for choice preferences when both targets appeared together, while the sequential condition allowed us to reveal how new information could alter decisions.

**Figure 2.**
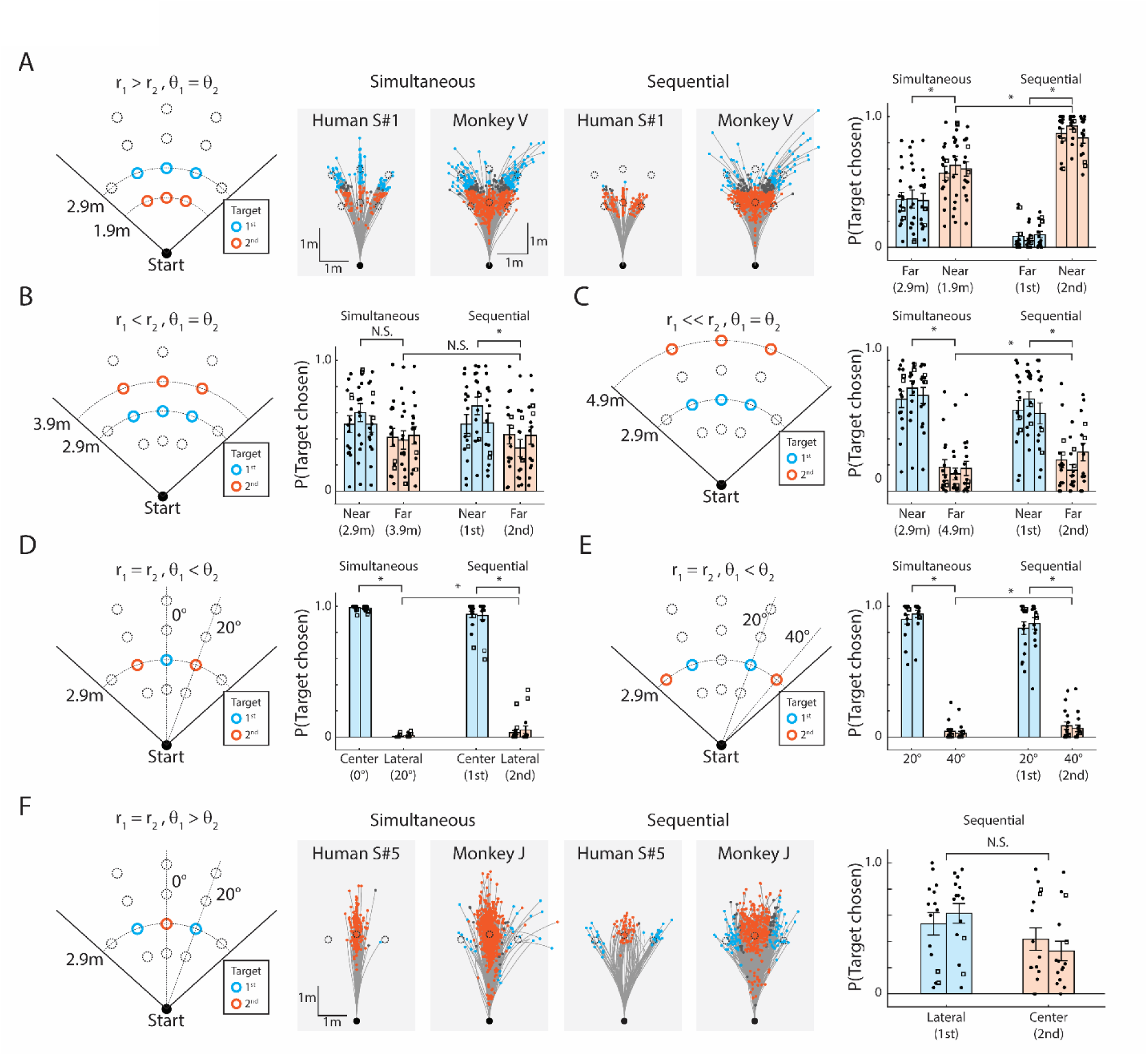
Target Selection Strategy under Simultaneous and Sequential presentations (A-C) Target preference for two targets at different distance and fixed angles: **(A)** Left: three target pairings (r₁ = 2.9 m; r₂ = 1.9 m). Middle: movement trajectories from representative human and monkey sessions for simultaneous and sequential presentations. Cyan dots indicate trials in which the first target was selected; orange dots indicate trials in which the second target was selected. Right: group-level target choice distributions. Bars are grouped by the lateral position of the first target (left, center, right). Filled circles denote human subjects; open rectangles denote monkey subjects. **(B–C)** Summary data for distance pairings of **(B)** r₁ = 2.9 m, r₂ = 3.9 m, and **(C)** r₁ = 2.9 m, r₂ = 4.9 m. **(D-F) Target preference for two targets at different angles and fixed distances:** **(D)** Targets positioned at θ₁ = 0°, θ₂ = ±20°; **(E)** θ₁ = ±20°, θ₂ = ±40°; **(F)** θ₁ = ±20°, θ₂ = 0°. Middle panels in **(F)** show representative trajectories from human and monkey sessions. Bars in **(D–F)** are grouped by the lateral side of the eccentric target (left, right). Error bars represent mean ± SEM. Statistical comparisons were conducted using paired t-tests (N = 15); *P < 0.05, N.S. for P > 0.05. Exact p-values are reported in the main text.

In trials where the two targets differed in distance but were positioned at the same angle (**Figure 2A**), both humans (N = 13) and monkeys (N = 2) consistently favored the closer target (r₁ = 1.9 m) over the farther target (r₂ = 2.9 m), whether the targets were presented simultaneously (P = 1.14 × 10⁻³, paired t-test, N = 15) or sequentially (P = 1.29 × 10⁻²⁶). This preference was significantly stronger in the sequential presentation – when the closer target appeared second (P = 2.56 × 10⁻¹²). Given that the second target was both more recent and required less effort to reach, subjects were more likely to select it, reflecting a sensitivity to both spatial cost and the timing of new information.

This recency modulation persisted even when both targets were farther away (**Figure 2B-C**). For modest distance differences (**Figure 2B**, r_1_ = 2.9 m < r_2_ = 3.9 m), the preference for the nearer target was weak in the simultaneous presentation (P = 0.0896) but became significant when the farther target was sequentially presented (P = 4.24 × 10^−2^). With a larger distance difference (**Figure 2C**, r_1_ = 2.9 m << r_2_ = 4.9 m), subjects consistently favored the nearer target in both simultaneous (P = 1.15 × 10^−11^) and sequential presentation (P = 1.31 × 10^−6^), yet subjects were still slightly more likely to select the farther, more recently presented target during sequential presentation (P = 4.47 × 10^−3^). These results suggest that subjects may favor a more recent target even when it incurs greater effort. Together, they indicate that proximity strongly guides navigational choice, while temporal recency modulates this preference— particularly when the associated effort cost is moderate.

We next examined trials in which the two targets differed in angular position (**Figure 2D-F**). Subjects strongly preferred less eccentric options. When one target was central and the other eccentric (θ₁ = 0°, θ₂ = ±20°, **Figure 2D**), central choices dominated in both simultaneous (P = 1.64 × 10⁻⁴⁴) and sequential (P = 6.07 × 10⁻²¹) trials. However, when the eccentric target appeared second, its selection increased slightly (P = 2.63 × 10⁻²), again reflecting the influence of recency. A similar pattern emerged when both targets were eccentric but differed in magnitude (**Figure 2E**; |θ₁| = 20°, |θ₂| = 40°): the less eccentric option was favored in both simultaneous (P = 1.03 × 10⁻²¹) and sequential (P = 2.44 × 10⁻¹⁵) conditions, yet the later-presented, more eccentric target was significantly more likely to be chosen (P = 2.88 × 10⁻⁴). These results mirror the distance-based patterns: subjects exhibit a robust bias toward central targets, but this preference can be modulated by the recent evidence.

The most dynamic behavior was observed when the first target was eccentric and the second was central (**Figure 2F**; θ₁ = ±20°, θ₂ = 0°). Subjects often initiated steering toward the initial, eccentric target but diverted their path after the appearance of the central target—indicating a change of mind.

This produced a roughly even distribution of final choices, with no significant preference between the two options (P = 0.0811). This behavior suggests a trade-off between recency and effort: while subjects tended to favor the more recently introduced target, doing so in this case required a substantial redirection of movement.

Collectively, these findings show that both humans and monkeys integrate spatial cost and temporal order when selecting between competing goals. Across conditions, subjects exhibited a strong preference for closer and more centrally located targets, with recent target onset systematically biasing decisions in the sequential condition. These results suggest that humans and monkeys dynamically update their belief in navigational decisions given new sensory information. We next examine how eye movements reflect these belief updates, focusing on sequential trials where changes of mind were most prevalent—specifically, those in which the second target was less eccentric than the first (**Figure 2F**; see **Supplementary Figure S2** for eye movement analyses under other conditions).

### Primate Gaze Tracks Future Target Selection and Reflects Changes of Belief during Sequential Target Presentation

We found that subjects’ eye movements continuously tracked their evolving belief about the intended target during sequential target presentation. Upon the appearance of the first target, subjects made a saccade to the flashed location and, during navigation, maintained gaze near its remembered position— reflected by a gradual convergence of horizontal eye position and a downward shift in vertical position (**Supplementary Figure S3** for detailed eye movement analysis), consistent with previous findings from single-target trials ^23^. However, we observed that gaze trajectories systematically diverged depending on the subject’s final choice. When subjects ultimately selected the first, eccentric target, the eye initially saccaded toward it and continued to track its location without shifting toward the second target (**Figure 3A**). In contrast, when subjects changed their decision to pursue the newly introduced central target, gaze redirected from the lateral position toward the center, aligning with the shift in steering behavior (**Figure 3B**). These examples suggest that eye movements reflect evolving internal decisions—not merely passive responses to visual input.

**Figure 3.**
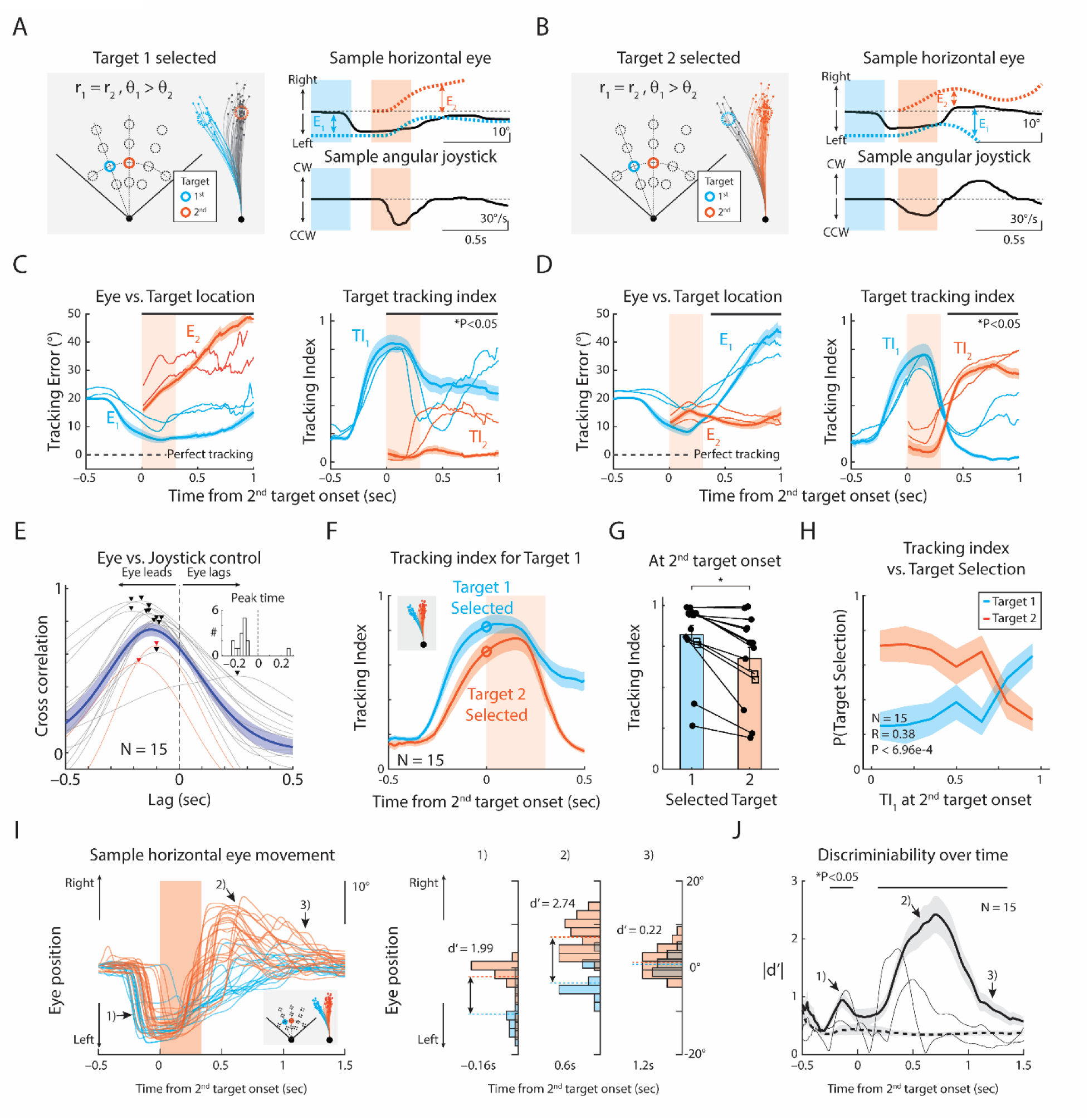
Eye Movements Reflect Target Selection and Changes of Mind during Sequential Target Presentation. **(A)** Left: movement trajectories from trials in which the subject selected target 1 (cyan, θ₁ = –20°) over target 2 (orange, θ₂ = 0°) during sequential presentation. Right: time course of horizontal eye position and angular joystick movement for an example trial. Dashed lines: ideal observer eye position for target 1 (cyan) or target 2 (orange). E₁ and E₂: tracking errors to each target center. **(B)** Same as **(A)**, but for trials in which the subject switched from target 1 to target 2. **(C)** Left: time course of target tracking error (E₁, cyan; E₂, orange), averaged across trials where target 1 was chosen, aligned to second target onset. Thick line: human average; thin lines: two monkeys. Right: average target tracking indices for target 1 (TI₁) and target 2 (TI₂). Black line above indicates significant differences between TI₁ and TI₂ (P < 0.05, paired t-test, N = 15). **(D)** Same as **(C)**, but for trials in which subjects switched from target 1 to target 2. **(E)** Cross-correlation between horizontal eye position and angular joystick movement. Triangles: peak lag (black: humans; red: monkeys). Inset: subject-level distribution. **(F)** Time course of TI₁, grouped by final target selection. Cyan: target 1; orange: target 2. **(G)** TI₁ at the onset of the second target, grouped by final target selection. TI₁ was significantly higher when subjects maintained their initial selection compared to trials in which they switched targets (*P < 0.05, paired t-test, N = 15). **(H)** TI₁ at second target onset versus target preference. TI₁ was positively correlated with the likelihood of maintaining the initial selection and negatively correlated with the likelihood of switching (Pearson’s correlation). **(I)** Left: horizontal eye position grouped by final choice. Right: discriminability index (d′) at three time points—(1) before, (2) after second target onset, and (3) near end of navigation. **(J)** Time course of d′ for humans (thick) and monkeys (thin), averaged across trials with θ₁ = ±20°, θ₂ = 0°. Dashed black line indicates control d′. Shaded area: SEM of human data. Black bar: time points where d’ is significantly greater than control (P < 0.05, paired t-test). Exact P-values are reported in the main text.

To quantify how accurately subjects tracked each target, we compared their eye movements to those of an ideal observer that continuously fixated the center of each target. Two key metrics were computed: (1) a tracking error (E₁ and E₂), defined as the deviation between the subject’s gaze and the ideal observer for each target; and (2) target tracking index (TI₁ and TI₂), a normalized measure ranging from 0 to 1 that reflects how closely gaze followed each target’s location (see Methods for details).

**Figures 3C** and **3D** compare these metrics separately: when subjects selected the initial, eccentric target (**Figure 3C**) and when they switched to the later, centrally located target (**Figure 3D**). In trials without a change of mind (**Figure 3C**), tracking error for the first target (E₁) remained consistently lower than that of the second target (E₂), and tracking index (TI₁) rose sharply following the first target’s appearance—indicating sustained, selective tracking of the initially chosen target. This pattern closely mirrors findings from single-target navigation tasks ^23^ and confirms that subjects actively tracked the intended goal location, rather than simply reacting to transient visual cues.

In contrast, when subjects changed their mind and selected the newly introduced central target (**Figure 3D**), eye movements exhibited a two-phase pattern: gaze initially tracked the first target, as before, but then redirected toward the new one. This behavioral shift was accompanied by a rapid reversal in tracking metrics. Around 350 ms after the second target appeared, tracking error E₁ increased and tracking index TI₁ declined, while E₂ dropped below E₁ and TI₂ rose markedly—coinciding with the shift in steering direction. These dynamics indicate that gaze tracks evolving internal decisions and reveals changes of mind during navigation, even after the target has disappeared from view (see **Supplementary Figure S2** for additional configurations). Collectively, these findings confirm that primates continuously track their intended—but invisible—goal, and that gaze trajectories reliably reflect belief updates in real time. However, this raises the question: do eye movements precede or merely follow motor action?

To quantify the temporal relationship between eye movements and motor actions, we computed the cross-correlation between horizontal eye position and angular joystick movement (**Figure 3E**). We found that eye movements consistently preceded steering responses by approximately 150 ms in most (12/13) humans and both monkeys. This lead time suggests that gaze and motor output are tightly linked but temporally dissociable—possibly reflecting parallel processing of sensory information and action planning ^26,27^.

Given this temporal correlation, we next asked whether early eye movements could predict subsequent changes of mind—particularly at the moment when the second target appeared (**Figure 3F**). We found that TI₁ measured at the onset of the second target, was significantly greater in trials where subjects maintained their initial choice compared to trials where they changed their decision (**Figure 3G**; P = 4.80 × 10⁻⁴, paired t-test, N = 15). Across subjects, TI₁ at the time of second target onset correlated positively with the probability of maintaining the initial selection and negatively with the likelihood of switching (**Figure 3H**; Pearson’s R = 0.38, P = 6.96 × 10⁻⁴, N = 15; see **Supplementary Figure S4** for correlations with final stop position).

We next asked: when do eye movements begin to predict a future change of mind? Given that the target-tracking index at the moment of the second target’s appearance predicted final choice, we examined whether gaze could reveal upcoming switches even earlier. In trials where subjects eventually changed their decision, horizontal eye position began to diverge from non-switch trials 200 ms before the second target appeared (**Figure 3I**, time point 1), suggesting that weakening belief in the initial target may precede reconsideration. After the second target appeared, gaze rapidly shifted toward the newly chosen target (time point 2), and later converged centrally as subjects approached the goal (time point 3). A time-resolved discriminability index (|d′|) confirmed this sequence: it rose significantly before second target onset, peaked shortly after, and diminished near the end of navigation (**Figure 3J**; P < 0.05, paired t-test vs. control, N = 15). These results suggest that changes of mind may arise from early uncertainty about the initially selected target.

Because gaze position prior to the second target onset diverged depending on the eventual decision, we asked whether the quality of gaze tracking—how precisely subjects followed the first target—reflected uncertainty in the initial representation. To test this hypothesis, we developed a model to estimate subjects’ internal estimate of the first target’s location prior to the second target’s appearance (**Figure 4A**). The model inferred each subject’s intended goal on a trial-by-trial basis using the seed state—defined as early gaze and movement trajectories before the second target onset (see Methods). To ensure that the model reflected commitment to the first target, it was trained only on trials where subjects ultimately chose it. We applied principal component analysis (PCA) to reduce the dimensionality of the seed state and trained a generalized linear model (GLM) to predict the final stopping position, used as a proxy for internal belief. The model accurately predicted both radial and angular coordinates of the stop position for both humans and monkeys (**Figures 4B–C**), confirming that early gaze and movement trajectories carry sufficient information to reconstruct belief about the initial target.

**Figure 4.**
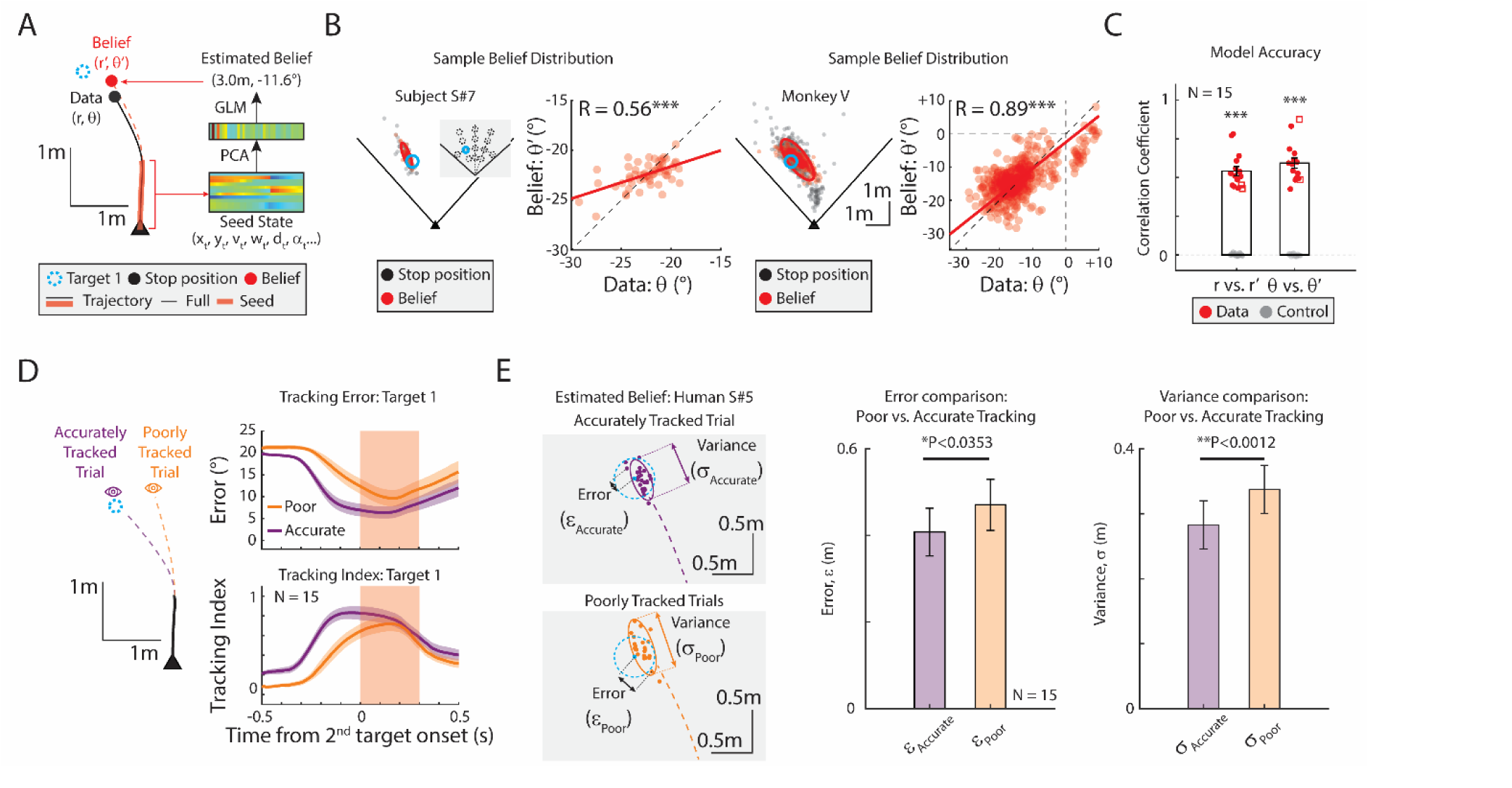
Belief Estimation Model and the Role of Gaze Accuracy in Belief Precision. **(A)** Belief estimation model. On each trial, internal belief about the first target location was estimated based on initial gaze position and trajectory before the second target appeared. Seed latent variables were extracted using PCA, and a GLM was trained to predict final stop positions (r′, θ′). **(B)** Beliefs of the first target (red) were compared with observed stop positions (black), for human (left) and monkey (right) subject. Belief of angular position, θ’, and ground truth angular position, θ, were significantly correlated, demonstrating that the model accurately predicts the belief. **(C)** Correlation coefficients for (r’ vs. r) and (θ’ vs. θ) were calculated for each subject. Belief estimation accuracy was significantly higher than the control estimation (gray) calculated from shuffled label. Filled circles: human individuals. Open rectangles: monkey subjects. **(D)** Trials were split by tracking performance into accurately tracked (purple) and poorly tracked (yellow) groups. Tracking error and tracking index were plotted relative to second target onset. **(E)** Belief precision by tracking accuracy. Belief distributions from each group were characterized by error (ε) and variance (σ) of fitted Gaussians (left). Poorly tracked trials had significantly less accurate predictions (middle; ε_Poor_ > ε_Accurate_, *P<0.05) and larger variance (right; σ_Poor_ > σ_Accurate_, **P<0.01, Wilcoxon sign rank test, N = 15), indicating higher uncertainty than accurately tracked trials. Error bars represent SEM, and statistical significance is denoted (*P<0.05, **P<0.01, ***P<0.001).

We then examined whether belief precision varied with gaze accuracy. Trials were split into two groups—accurate vs. poor tracking of the first target—based on target-tracking metrics, and their predicted belief distributions were compared (**Figure 4D**). In accurately tracked trials, belief estimates were tightly clustered around the true target, indicating high certainty. In contrast, poorly tracked trials yielded belief distributions with larger errors (ε) and greater variance (σ), reflecting higher uncertainty. Both prediction error (P = 3.53 × 10⁻^2^) and belief variance (P = 1.16 × 10⁻³) were significantly higher in poorly tracked trials compared to accurately tracked trials (Wilcoxon signed-rank test, N = 15). These results demonstrate that imprecise gaze is linked to uncertain belief representations, supporting the idea that early uncertainty about the initial target underlies subsequent changes of mind. Taken together, these results demonstrate that primate eye gaze dynamically reflects evolving internal beliefs and predicts changes of mind during sequential decision-making. Next, we examine how gaze dynamics integrate with other task variables to shape future navigational behavior.

### Eye Movement Conveys Task-Related Information for Naturalistic Navigation

To investigate whether latent variables during navigation encode upcoming choices, we examined key features of subjects’ internal state—including velocity, heading, distance and angle to both targets, and gaze position. These variables evolve continuously and interact dynamically, forming a closed-loop system between perception and action (**Figure 5A**).

**Figure 5.**
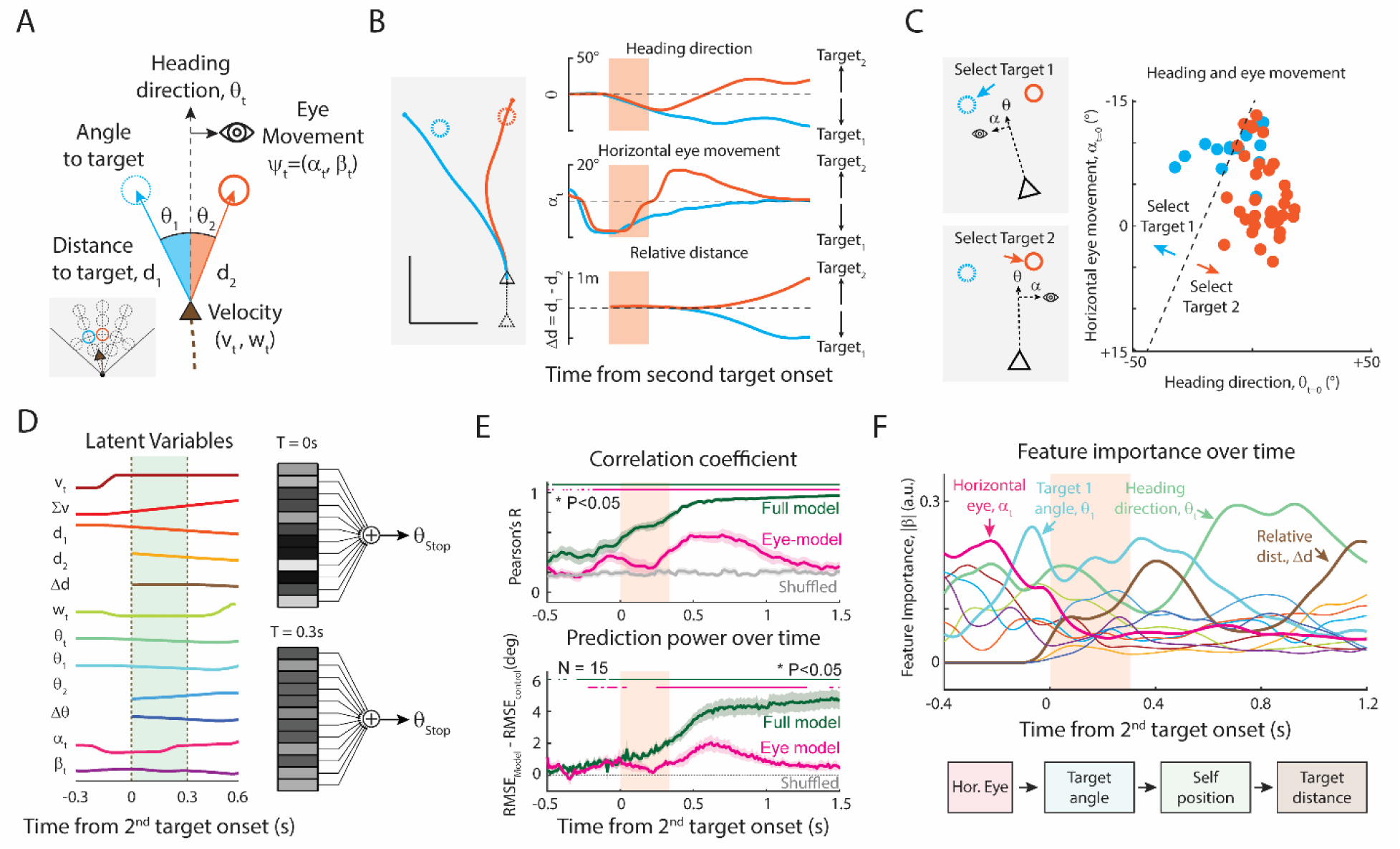
Eye Movements Convey Latent Information for Future Target Selection. **(A)** Latent variables used to model navigational decisions over time. Linear velocity (vₜ), angular velocity (wₜ), distance to target 1 and 2 (d₁, d₂), relative angle to target 1 and 2 (θ₁, θ₂), heading direction (θₜ), and horizontal and vertical eye position (αₜ, βₜ) were computed at each time point. **(B)** Example time courses of latent variables aligned to the onset of the second target. Trials are grouped by final target selection: cyan lines indicate trials where target 1 was selected; orange lines indicate trials where target 2 was selected. **(C)** Example classification of final target selection using heading direction (θₜ) and horizontal eye position (αₜ) at the time of second target onset. Cyan and orange dots indicate final target choices; black dashed line denotes the classification boundary. **(D)** Linear regression models predicting final stop position (θ_stop_) from latent variables. **(E)** Model performance over time. Top: Pearson’s correlation coefficient between predicted and observed θ_stop_; Bottom: Prediction power, defined as RMSE_Model_ – RMSE_Control_ (RMSE = Root-mean-squared-error). Green: full model using all latent variables; magenta: horizontal eye position only; gray: control model using shuffled labels. Colored bars above indicate time points with significant prediction performance relative to the control model (P < 0.05, paired t-test). **(F)** Feature importance over time in the full model. Average absolute value of regression coefficients (|β|) for each feature is shown, highlighting the relative contribution of each variable across time. See **Supplementary Figure S5** for full details.

Distinct patterns of latent variables emerged when comparing trials based on the final choice. Although subjects occupied similar positions at the onset of the second target, their heading direction, horizontal eye position, and relative distance rapidly diverged in a choice-specific manner. For instance, trials ending in selection of the initial (eccentric) target were characterized by leftward heading and gaze, while those culminating in a switch to the second (central) target exhibited rightward deflections (**Figure 5B**). These differences reflected consistent motor and gaze signatures associated with distinct navigational decisions.

To illustrate this, we trained a simple linear classifier using just two variables—heading direction and horizontal eye position—measured at the onset of the second target. This minimal model successfully predicted final target selection with high accuracy (**Figure 5C**), indicating that gaze and heading already encode the subject’s evolving internal belief at the time of the new evidence.

For a comprehensive analysis, we quantified 12 latent variables: linear and angular velocities (v_t_, w_t_), cumulative distance travelled (Σv_t_), horizontal and vertical eye movements (α_t_, β_t_), heading direction (θ_t_), linear distance to each target (d_1t_, d_2t_), relative linear distance (Δd_t_), angular distance to each target (θ_1t,_ θ_2t_), and relative angle (Δθ_t_). Each variable was aligned to the second target’s onset, and a linear regression model was used at each time frame to predict each trial’s final stop position θs_top_, on a trial-by-trial basis (**Figure 5D**). Model performance was assessed by Pearson’s correlation and root-mean-squared-error (RMSE) between the predicted and observed θs_top_.

As expected, the full model using all 12 variables yielded accurate predictions of θs_top_, while a control model using shuffled trial labels failed to predict. To isolate the specific contribution of eye movement to future planning, we focused our subsequent analysis on a simplified model using only horizontal eye position (**Figure 5E**). The temporal profile of model accuracy revealed two notable features. First, final stopping positions could be predicted even before the second target appeared (∼200 ms in advance). Second, a model using only horizontal eye position showed dual peaks in accuracy—one before and one after second target onset. The early peak reflected whether the subject had reliably tracked the first target (see **Figure 3F**); the later peak captures whether gaze had shifted to the new target, highlighting the role of eye movements in both maintaining and updating internal beliefs (**Figure 5E** and **Supplementary Figure S5**).

Finally, we assessed the temporal contribution of each latent variable by examining the absolute value of its regression coefficient (|β|; **Figure 5F**). Prior to the second target’s introduction, horizontal eye position was the most informative feature for predicting stopping behavior—consistent with **Figures 3F** and **5E**. This suggests that gaze reflects the strength of the belief about the first target and influences whether that belief is maintained. After the second target onset, heading direction and relative distance jointly became the dominant predictors, reflecting rapid integration of new sensory input. These findings indicate that eye movements not only track task-relevant latent variables but also convey belief-related information that guides flexible navigation.

## Discussion

Our study explored the role of eye movements in shaping future decisions, focusing on how humans and monkeys flexibly navigate toward one of two transient targets in a dynamic virtual environment. By presenting targets either simultaneously or in sequence, we showed that new sensory input could revise ongoing behavior, prompting subjects to update their initial motor plan. Eye movements were closely coupled to final target selection, continuously tracking the intended goal and adapting in parallel with shifts in navigational strategy during change-of-mind events. Through quantitative modeling, we demonstrated that gaze trajectories encoded predictive information about future actions, highlighting their critical role in dynamic decision-making under uncertainty.

### Adaptive Decision-Making in Dynamic Environments

The ‘firefly’ experiment, which requires path integration within a closed action-perception loop, challenges subjects to navigate toward a remembered goal by integrating optic flow. Previous research has established that successful navigation toward a remembered goal ^28,29^ depends on tracking the invisible target with their eyes ^23^. Variations of the firefly experiment have revealed that primates navigate effectively regardless of sensory modality ^30^, adaptively adjust their beliefs when self-motion or action space is perturbed ^21^, and even infer the moving goal’s location ^31^.

Building on this previous research, we developed an immersive virtual reality setting identical for both humans and monkeys, offering more naturalistic conditions to fairly compare species behaviors. We found a large consensus between human and monkey behavior, not only in task performance (**Figure 2**, **Supplementary Figure S1**) but also in the eye movement patterns and decision dynamics (**Figure 3**, **Supplementary Figure S2**). Although monkeys showed a minor tendency to favor recently introduced targets more than humans (**Figure 2C-D**)— possibly due to species-specific weighting of recent sensory evidence or differences in working memory capacity ^32^, both species employed similar strategies to adapt and plan in dynamic environments.

This robust inter-species similarity provided the ideal foundation to investigate the dynamic nature of decision revision. To this end, our sequential target paradigm extended the closed-loop design by introducing a second, equally rewarded target with different action costs and lower uncertainty. This approach created opportunities for subjects to revise their initial plans in response to new information. Because the targets appeared only briefly and required rapid integration (within a few hundred milliseconds), the task probed how subjects balance action cost against memory reliability. Our behavioral results revealed two consistent patterns (**Figure 2**): (1) both species preferred targets that minimized effort—closer targets requiring less forward movement or central targets requiring smaller turns; and (2) newly introduced targets were more likely to be selected, reflecting their lower memory demands and reduced uncertainty. These patterns are consistent with the idea that longer path integration degrades memory precision, making newer targets more reliable. This also aligns with prior findings showing that primates prefer more certain options—particularly when competing choices involve degraded or less reliable memory representations ^33^.

While change of mind has been widely investigated within the classical perceptual decision-making framework ^34–37^, previous research has mainly focused on confidence and decisions in open-loop conditions, where action does not influence perception. Amid growing interest in employing more naturalistic and continuous experimental paradigms to explore the brain’s cognitive capabilities ^10,12,38,39^, our findings help bridge the gap between traditional decision-making and continuous action selection ^11^ by illustrating how subjects dynamically revise their choices in response to new sensory input within a closed-loop environment.

### Role of eye movement in active sensory integration and future planning

Our study highlights the multifaceted role of eye movements in navigating uncertain and dynamically changing environments. Prior research has suggested that eye movements play a crucial role in various processes, including attention allocation ^40,41^, evidence accumulation ^42,43^, information retrieval ^44^; for a review, see ^8,45^. Extending these findings into an active, closed-loop context, our results demonstrate that both human and monkey eye movements continuously integrate task-relevant sensory evidence during goal-directed navigation ^9,46,47^.

As depicted in **Figure 3** and **Supplementary Figures S2-3**, primates’ eyes rapidly saccaded to newly introduced targets and continued to track the selected target throughout navigation. This behavior suggests that gaze is not merely reactive but contributes actively to latent computations by integrating optic flow and monitoring key task variables, such as the sudden appearance of a new goal.

Our findings elucidate the significant role of eye movements in navigating uncertain environments, particularly emphasizing how accurately tracking an invisible target reflects the robustness of internal beliefs. Recent work^25^ demonstrated that gaze during navigation embodies the navigator’s evolving internal beliefs about goal locations, rather than simply mirroring external stimuli. Consistent with that view, we found that accurate tracking was associated with more stable decisions, while degraded tracking predicted an increased likelihood of switching to the second target (**Figure 3F-H**). These results suggest that gaze precision serves as an implicit measure of belief certainty: high precision indicates strong commitment to the initial target, while poor precision reflects uncertainty, increasing the probability of a change of mind when new evidence appears (**Figure 4**). Thus, gaze is not merely a passive readout of a committed decision but a dynamic signal that evolves with—and reveals— the decision-making process itself.

**Figure 5** further supports this interpretation by quantifying the predictive value of gaze for final navigational decisions. The model shows that trial-by-trial gaze features carry sufficient information to predict the eventual decision outcome, particularly before overt motor actions are initiated (**Figures 3F and 5**). Together, these findings position eye movements as a predictive correlate of belief updates during active navigation, serving both as a sensory-integration mechanism and as an early indicator of upcoming decision.

## Methods

### Subjects

The experiments involved two rhesus macaques and thirteen human participants. All human subjects were over 18 years old, with six males included. Prior to participating in the experiment, the human subjects were unaware of the study’s purpose and provided their informed consent.

### Experimental setup

Participants were tasked with virtually navigating to and stopping at the location of a briefly presented target (i.e., the “firefly”). Participants used a two-degree-of-freedom analog joystick (linear and angular velocity; M20U9T-N82, CTI Electronics) to navigate the virtual environment. Visual stimuli were rendered via custom code in Unity (programmed in C#) and displayed on a virtual reality (VR) head-mount display with a built-in eye tracker at a frame rate of 90Hz. The subjective vantage point was set to a height of 10 cm above the ground, and participants had a field of view of 110.1° of visual angle. The virtual world consisted of a circular ground plane, where textural elements and targets appeared. Each textural element was an isosceles triangle (8.5 x 18.5 cm²) randomly repositioned and reoriented within the ground plane and had a limited lifetime of 300 ms to avoid serving as landmarks. The density of the elements was set at 50 elements per square meter.

On each trial, subjects were re-positioned at the center of the virtual world, and the target (black circle of radius 25 cm, luminance and color matched to ground plane triangles) was displayed. On 10% of trials, only a single target was presented for 300 ms, in one of 14 predefined positions which varied in radial distance (190 cm, 290 cm, 390 cm, and 490 cm) and angle (−40°, −20°, 0°, 20°, and 40°). By design, the closest distance between targets was 100 cm. On 45% of trials, two targets appeared simultaneously in 15 different combinations of positions (**Figure 1B**, **Supplementary Figure S1**). On the remaining 45% of trials, the two targets were sequentially presented using the same 15 pair combinations, but with random inter-target-interval (uniform random distribution, minimum = 200 ms, maximum = 550 ms). These three conditions were randomly interleaved throughout the session. Because our primary objective was to investigate the continuous updating of an ongoing navigational plan, we excluded trials where subjects initiated movement only after the second target was introduced (Exclusion percentage = 17 ± 10 %). This criterion ensured that we analyzed decisions made based on pre-movement belief, isolating planning rather than reactive corrections. We additionally confirmed that this exclusion does not affect the main result of our analysis.

Human subjects received graded feedback based on their accuracy in reaching a target. For the 2-target trials, two points were awarded if stopping within 25 cm of any target, one point for stopping within 25 cm to 50 cm, and no points otherwise. Participants received compensation based on their score. The maximum linear speed varied randomly between 2 m/s and 3 m/s for each trial, while the angular speed remained fixed at 90°/s. Before the first session, all human participants underwent 40 practice trials (20 trials for single target, and 20 trials for two targets) to familiarize themselves with joystick movements and the task structure. Each human subject completed at least 2800 trials across multiple sessions (typically 4 sessions), which yields (on average) 20 trials per single target position and (on average) 84 trials per two target combination.

The monkey subjects also received a juice reward based on the distance between its stop position and one of the targets. A large amount of juice was given if it stopped within 25 cm of any target, a small amount of juice was given if it stopped within 25 cm to 50 cm, and no juice was given otherwise. The monkeys were previously trained to catch a single target, but never been exposed to experiment with two targets. All other experimental setups remained the same as those for the human subjects. The monkey completed at least 8000 trials across multiple sessions (approximately 15 sessions across several days), providing enough trials for each condition.

### Eye tracking

For human subjects, at the beginning of each block of trials, participants calibrated the VIVE Pro eye tracker using the inbuilt Tobii software. The calibration process involved fixating on several points displayed in the VR environment. Both eyes were tracked, and the participant’s point of foveation (x-y coordinates), object of foveation (ground, obstacles, boundaries, etc.), pupil dilation, eye openness, and other variables of interest were recorded on each frame using the inbuilt software.

For the monkey, a VR headset (HTC Vive) with a mounted 200 Hz eye-tracker add-on (Pupil Labs, Berlin, Germany) was secured on the chair in front of the monkey to display visual stimulus. The headset and eye-tracker were manually modified to account for the smaller interpupillary distance of monkeys. The calibration process involved fixating on 9 points displayed in the VR environment. The same variables, except pupil dilation, were recorded on each frame using the inbuilt software.

### Psychometric analyses

We compared the performance between the presentation conditions as follows. For each subject and each condition, we first constructed a psychometric function by calculating the proportion of correct trials as a function of (hypothetical) reward boundary (**Supplementary Figure S1D-E**). We used single target condition as a control, to set the baseline of each subject’s performance. From the obtained ROC curve by plotting the proportion of correct trials for each value of hypothetical reward boundary, we used the area under this ROC curve to obtain an accuracy measure.

### Eye gaze analyses

To simplify the analysis, we represented the subject’s eye gaze position using two standard degrees of freedom: (i) Horizontal eye movements, which was quantified as the average lateral position of the two eyes, *α* = (*α_l_* + *α_r_*)/2, and (ii) Vertical eye movements, which was quantified as the average vertical position of the two eyes, *β* = (*β_l_* + *β_r_*)/2, following the previous reference ^23^.

### Modeling the ideal observer

To evaluate whether the subject’s eye gaze was following the target, we formulated an ideal observer that consistently tracked the center of the (invisible) target. The location of the target relative to the midpoint of the subject’s eyes at time t is denoted as (x_t_, y_t_, z_t_). The ideal lateral displacement of the eye 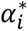 was calculated as 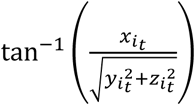, and the vertical displacement of the eye 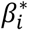 was calculated as 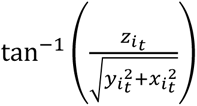. In our environment, z_t_ is determined by the camera height (10 cm) and remains constant, while x_t_ and y_t_ change continuously as the subject moves toward the target i.

### Target tracking error and target tracking index

To compare the observed eye movement with the ideal observer, we calculated the target tracking error between the observed eye movement *φ_t_* = (*α_t_*, *β_t_*), and the ideal observer that tracks either target 1, 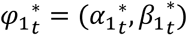 or target 2, 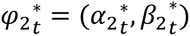. The target tracking error was computed as follows:

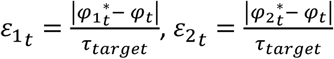

where | | denotes the length of the vector, and *τ* _target_ denotes the span of the target at time t.

Using this target tracking error, we quantify the ‘target-tracking index’ of observed eye movement by following:

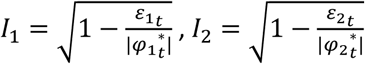

A value of 1 (i.e., *ε_t_* = 0) corresponds to ideal tracking of the target, and a value near 0 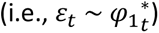 corresponds to not tracking the target. A few features were additionally accounted: 1) When the magnitude of the error is larger than the ideal eye movement, we clipped the target tracking index to zero, because it is essentially not tracking the target, 2) When ideal observation is impossible to make (i.e., when subject traveled beyond the target), the target tracking index was nullified, and 3) target tracking index was not calculated from 100 ms before the subject stopped their navigation until the end of the trial to eliminate the effect of stop.

### Defining the selected target

The location of the targets and subjects’ endpoint were expressed in polar coordinates, with a radial distance (target = r_1_, r_2_ and response = *r̅*) and eccentricity from straight ahead (θ = θ_1_, θ_2_ and *θ̅*). We excluded trials where (1) the subject stopped within 0.5 m of the origin (i.e., made essentially no movement), or (2) the subject failed to stop at all (max duration exceeded). For the remaining trials, target choice was defined as follows: if the subject’s stop point was within 0.5 m of one of the targets, that target was counted as chosen. If the stop point wasn’t within 0.5 m of either target (no reward), we assigned the choice to the target which the subject stopped closer to – as long as it was at least 33 cm closer than the other target. The 33 cm threshold corresponds to one-third of the minimum inter-target distance (100 cm), partitioning the inter-target space into three equal zones (target 1, target 2, indecisive). If distances to both targets were within 33 cm, we next compared angular error, and the target with a significantly smaller eccentricity error (with a margin of 2°) was defined as a selected target. If neither distance nor angular cues clearly indicated one target (within 33 cm and 2°), the trial was labeled “indecisive” and excluded from choice analyses (4.4 ± 2.4% of trials).

### Cross-correlation between eye and joystick movement

We computed eye–joystick cross-correlations for sequential-target trials (focusing on the configuration where the first target was lateral and the second target was central; see **Figure 3**). We treated horizontal (lateral) eye position and angular joystick velocity as one pair of signals, and vertical eye position and linear joystick velocity as another pair. We calculated the cross correlation between the lateral eye movement, α_t_ and angular joystick movement, w_t_ from trial onset until the subject stopped.

### Time-resolved discriminability index (d’)

To assess the capacity of horizontal eye movements to differentiate between target selections over time, we calculated the discriminability index, d’, for each subject by the following equation:

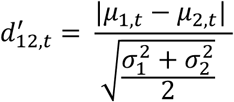

where *μ*_1,*t*_ and *μ*_2,*t*_ denotes mean of the horizontal eye movements aligned with the second target onset, and *σ*_1_ and *σ*_2_ denotes standard deviations of these eye movements. To calculate the statistical significance, we calculated the surrogate d’ with the shuffled trial label and plotted against original d’ (**Figure 3J**) and measured the paired t-test between the original d’ and surrogate d’.

### Belief estimation model

To estimate the belief of the target position, we trained a PCA-GLM model using subjects’ latent state variables in the 500 ms before the second target appeared (**Figure 4**). We first selected a subset of trials in which subjects selected the first target as a final decision during sequential presentations. From this subset, we extracted linear velocity (v_t_), angular velocity (w_t_), heading direction (θ_t_), relative distance to the first target (d_1t_), relative angles to the first target (θ_1t_), current subject position (x_t,self_, y_t,self_, r_t,self_, θ_t,self_), and horizontal and vertical eye movements (α_t_, β_t_). All time-series latent variables were aligned to the onset of the second target, and a 500 ms window of data before the second target’s onset was used for prediction. To reduce the dimensionality of the data, principal component analysis (PCA) was applied, retaining components that explained 95% of the variance. These reduced latent variables were then used as inputs to a generalized linear model (GLM) to predict the final stop position (r_stop_, θ_stop_)—a proxy for subjects’ belief about the target location.

Model performance was validated by comparing predicted stop positions (r’, θ’) with observed stop positions (r_stop_, θ_stop_) in each trial. Pearson’s correlation coefficients were calculated to assess the accuracy of predictions for both radius (r’) and angular position (θ’). A control correlation was calculated using shuffled trial labels to ensure predictions were not due to chance. (**Figure 4A-C**)

To investigate the impact of tracking accuracy on belief, trials were categorized into accurately tracked and poorly tracked groups based on target tracking error. Trials with tracking errors exceeding the average error were classified as poorly tracked, while those with lower errors were categorized as accurately tracked. Belief distributions were quantified using two parameters: (1) error (ε), the distance between the predicted position (r’, θ’) and the first target position, and (2) variance (σ), computed as the larger eigenvalue of a fitted Gaussian mixture. (**Figure 4D-E**)

### Predicting stop position from latent variables

To assess which latent variables convey the most information for future navigation, we identified a set of 12 variables that capture the dynamic state of a subject within the navigation task. These variables include linear velocity (v_t_), cumulative distance traveled (∑v_t_), distances to the first and second targets (d_1_, d_2_), the difference in distances to the two targets (Δd), angular velocity (w_t_), heading direction (θ_t_), angles to the first and second targets (θ_1_, θ_2_), the change in angular distance between the targets (Δθ), and horizontal and vertical eye movements (α_t_, β_t_), all aligned to the onset of the second target (**Figure 5**).

We employed Lasso regression (Least Absolute Shrinkage and Selection Operator) model to predict the final stopping position θ_stop_ using the 12 latent variables. The regularization strength of the Lasso was determined using cross-validation to identify the optimal lambda (λ) value that minimizes prediction error with 10-fold validation. We compared three Lasso models, 1) full model which used all 12 latent variables, 2) an eye-only model using only horizontal eye movement, and 3) control model which used all 12 variables but with shuffled trial label. To assess the model prediction power, we calculated 1) the Pearson’s correlation coefficient between the observed θ_stop_ and predicted θ_stop_, and 2) the root-mean-squared-error (RMSE) between the two (**Figure 5E**, see also **Supplementary Figure S5**). For the full model, we examined the temporal dynamics of the magnitude of Lasso regression model’s coefficients, |β|, which indicates the varying influence of each latent variable throughout the navigation task (**Figure 5F**). We also replicated these model predictions, using the variables aligned to trial onset (**Supplementary Figure S5D-F**).

### Statistical analysis

Group-level analyses treated each subject as one observation, pooling the 13 human participants and the 2 macaques (N = 15). Comparisons between conditions used two-tailed paired t-tests, except for Fig. 4E, which used the two-tailed Wilcoxon signed-rank test. Correlations were quantified with Pearson’s correlation coefficient. Error bars denote s.e.m. across subjects.

## Ethics Declarations

All human experimental procedures were approved by the Institutional Review Board (IRB) at New York University (IRB-FY2019-2599). Prior to participating in the experiment, human subjects were provided their informed consent. All animal care and experimental procedures were approved by the Institutional Animal Care and Use Committee (IACUC) at New York University (Protocol 18-1499).

## Author Contributions

W.C., P.A., and D.E.A. conceptualized the study. W.C., P.A., and D.E.A. developed the methodology. W.C., P.A., G.A., and D.E.A. validated the experimental paradigms. W.C. visualized the data. W.C. and D.E.A. conducted the formal analysis. W.C. and D.E.A. performed the investigation. W.C. and G.A. curated the data. W.C. wrote the original draft. W.C. and D.E.A. reviewed and edited the manuscript. D.E.A. supervised the study, administered the project, and acquired funding.

## Acknowledgements

We thank Tim Haoran Ding for his technical assistance with software development, and Akis Stavropoulos for helpful discussions and his contribution to the development of the VR headset. This research was supported by KBRI basic research program through Korea Brain Research Institute funded by Ministry of Science and ICT (26-BR-03-01 and 26-BR-ISD-02, to W.C.), and by NIH NINDS U19 grant NS118246 and R01 grant NS127122 (to D.E.A.).

## Data Availability

The data supporting this study will be made publicly available in an OSF repository upon publication.

## Code Availability

The code used for analysis and modeling will be made publicly available in an OSF repository upon publication.

---

**Supplementary Figure 1.**
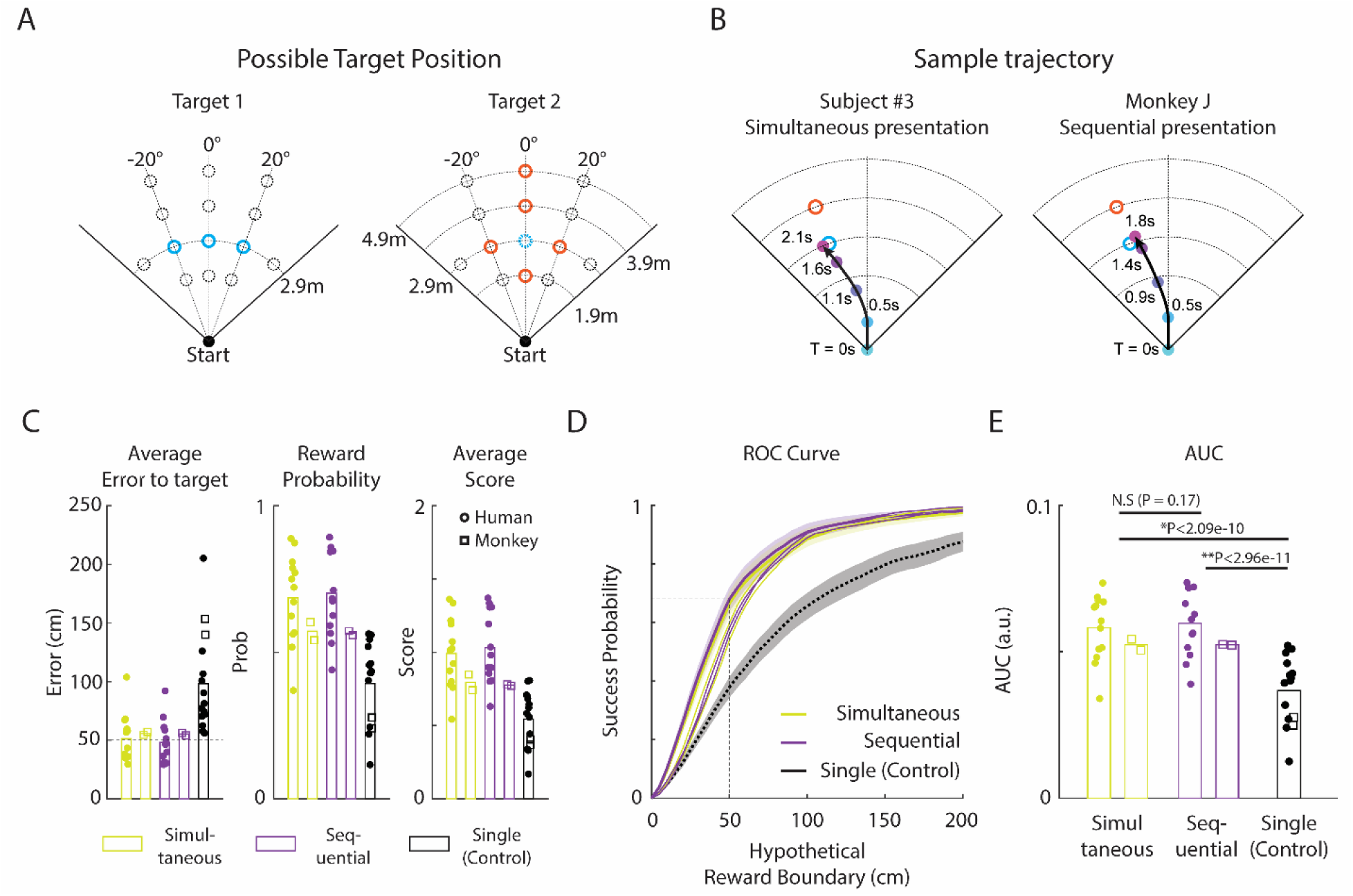
Possible Combinations of Target Presentation, Exemplary Trajectories, and Performance Comparison between Simultaneous and Sequential Presentation, related to Figure 1. **(A) Possible target positions**. The first target may appear at one of three positions with a 2.9 m radius at −20°, 0°, or +20° angles. The second target may appear in one of five positions: 1) the same angle but at 1.9 m, 2) 3.9 m, 3) 4.9 m, 4) the same distance but at −20°, or 5) +20° relative to the first target. **(B) Sample trajectories** of a human participant and monkey J during simultaneous and sequential presentations, respectively. Cyan and orange circles denote the first and second target positions, respectively, with black lines indicating sample trial trajectories. **(C) Performance of human and monkey subjects** under simultaneous or sequential presentation conditions. Error (left) presents the distance between the stop position vs. nearest target, averaged across all trials (rewarded and unrewarded). Dashed line shows the reward boundary (50 cm). Reward probability (middle) presents the ratio of trial with error < 50 cm. Score (right) is calculated as follows: 2 points are awarded for each trial if the error is less than 25 cm, and 1 point is given for trials where the error is between 25 cm and 50 cm. Yellow and purple dots and bars denote simultaneous and sequential presentation conditions, respectively, while black dots and bars indicate single target presentation. Filled circles represent human subjects and open rectangles represent the two monkey subjects. Humans and monkeys exhibited similar trends in all aspects, with performance in simultaneous and sequential presentations being comparable and superior to the single target condition. **(D) Receiver operating characteristic (ROC) example**, computed using a variable hypothetical reward boundary (Methods). Thick lines: Human data; ROC curve for simultaneous (yellow) and sequential (purple) conditions, with the black line representing the single target condition. Thin lines: data from two monkeys. **(E) Summary area-under-curve (AUC)** in the three conditions. Simultaneous and sequential presentations showed no significant difference in AUC performance (N.S., P = 0.17), but were significantly better than the single target conditions (*P < 2.09 × 10^−10^, ** P < 2.96 × 10^−11^, for simultaneous vs. single and sequential vs. single, respectively. Paired t-test, N = 15; 13 humans and 2 monkeys).

**Supplementary Figure 2.**
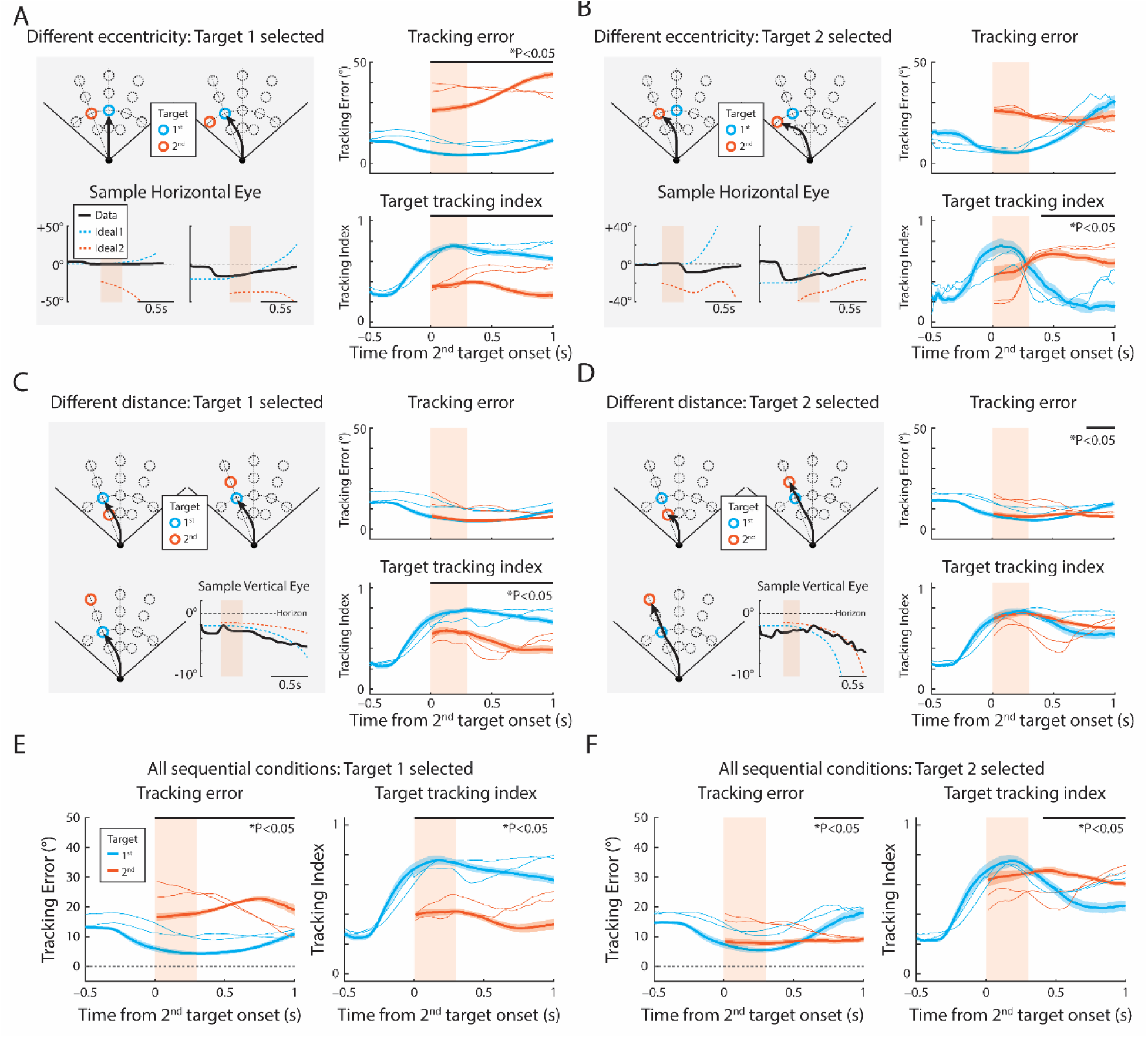
Target Tracking Error and Target Tracking Index under all Sequential Presentation Conditions, related to Figure 3. **(A-B)** Target tracking error and target tracking index when the second target is presented more laterally (e.g., (θ_1_, θ_2_) = (0°, ±20°) and (θ_1_, θ_2_) = (±20°, ±40°)). **(A)** When subjects select the first, less-laterally positioned target, the horizontal eye movement follows the first target position (left; cyan dotted line) rather than the second target (orange dotted line). The right panel shows that tracking error E_1_ is significantly smaller than E_2_ (top) and tracking index TI_1_ is significantly higher than TI_2_ (bottom) for all time points. **(B)** When subjects select the second, eccentric target, the horizontal eye movement shifts toward the more lateral target (bottom left; orange dotted line). Tracking error E_2_ decreases over time (top right), and the target tracking index TI_2_ increases significantly approximately 390 ms after the second target onset. **(C-D)** Target tracking error and target tracking index when the second target is presented at the same angle but different distance (e.g., θ = θ_1_ =θ_2_, r_2_= 1.9 m or 3.9 m or 4.9 m). **(C)** When subjects select the first target at 2.9 m, the vertical eye movement follows target 1 (bottom left; cyan dotted line) rather than target 2 (orange dotted line). The tracking error E1 and E2 were not significantly different, but the target tracking index shows TI_1_ was significantly higher than TI_2_ for all time points. **(D)** When subjects select the second target with a different distance, the vertical eye movement follows target 2 (bottom left; orange dotted line) rather than target 1 (cyan dotted line). Target tracking error (top right) E_2_ becomes significantly smaller ∼780 ms after target 2 presentation, and target tracking index TI_2_ becomes subtly larger than TI_1_ although not significant, presumably due to low variance in vertical eye movement. **(E-F)** Target tracking error and tracking indices for all sequentially presented trials, when **(E)** subject selects target 1 and **(F)** subject selects target 2. **(E)** When a subject chose target 1, E_1_ is significantly smaller than E_2_, and TI_1_ is significantly higher than TI_2_ at all time point. **(F)** When subject chose target 2, E_2_ becomes significantly higher than E_1_ approximately 660 ms after target 2 onset, and target tracking index TI_2_ becomes significantly higher than TI_1_ approximately 400 ms after target 2 onset. Shaded area denotes the standard error of human subjects, and thin lines denotes the monkey representation. All paired t-tests were conducted with N = 15.

**Supplementary Figure 3.**
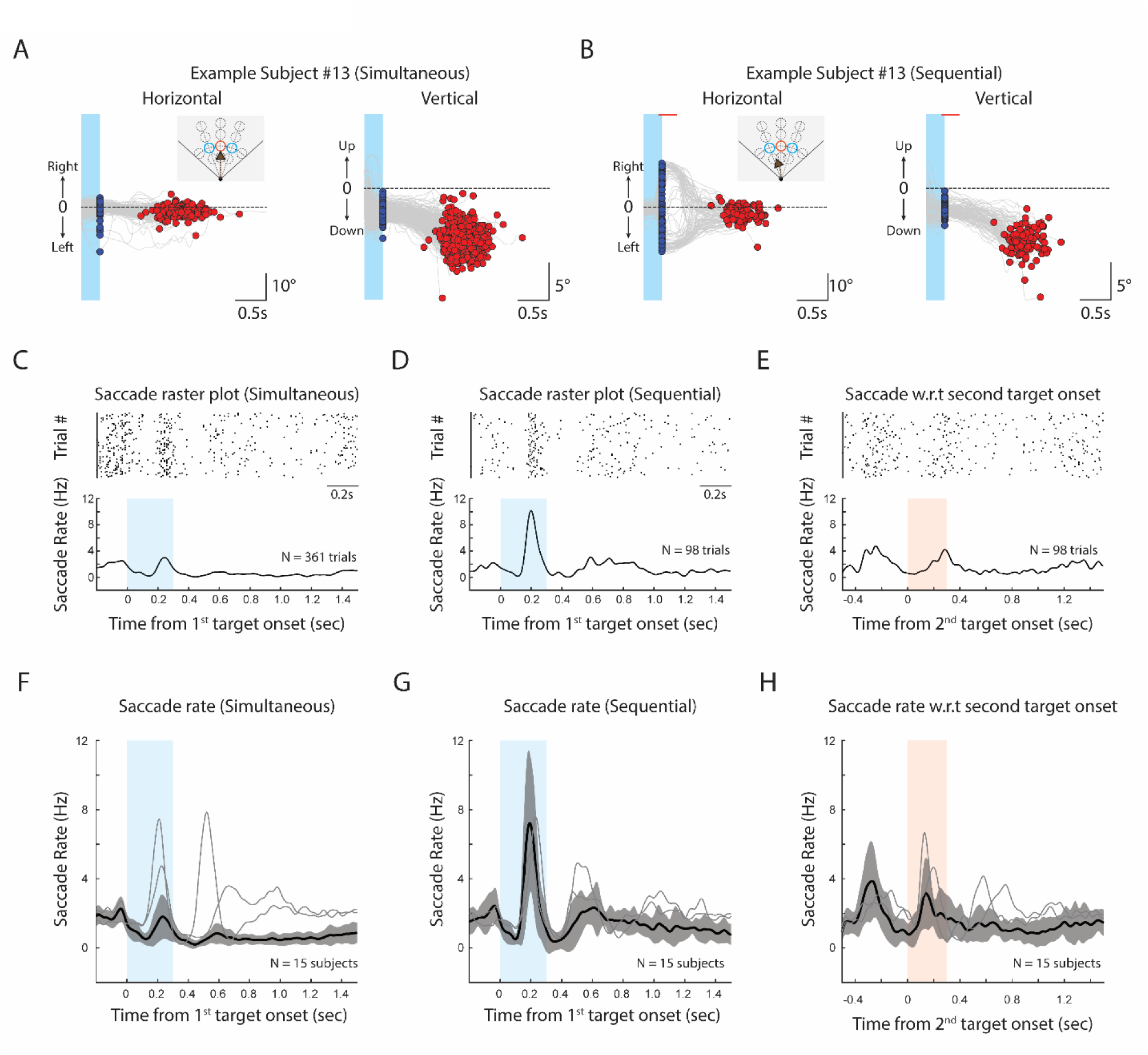
Eye movements during the task, related to Figure 3. **(A, B)** Sample human subject’s horizontal and vertical eye movements during **(A)** simultaneous and **(B**) sequential presentations when the two targets are positioned at the same distance but different eccentricity (Figure 2F). Blue dots denote gaze position when the target disappears, and red dots denote the final gaze position 100 ms before the subject stopped. **(A)** During simultaneous presentation, because the subject mostly chooses the central target, horizontal eye position does not make noticeable shift, but vertical eye position shifts downwards as the subject gets closer to the target. **(B)** During sequential presentation, because the target appears eccentrically, subjects make a lateral saccade, and their gaze shifts centrally as subject navigate, as shown previously ^23^. **(C, D)** Sample saccade raster plot and saccade rate aligned to 1^st^ target onset, during **(C)** simultaneous and **(D)** sequential presentation, in trials of panel A and B. Compared to the simultaneous condition, the first eccentric target during sequential presentation induces significantly more frequent and consistent saccades. **(E)** Sample raster plot and saccade rate aligned to the second target onset during sequential presentation. **(F-H)** Average saccade rate during **(F)** simultaneous, **(G)** sequential presentation, aligned to 1^st^ target onset, or **(H)** aligned to 2^nd^ target onset, respectively. Thick line: average saccade rate of 13 human subjects; Thin lines: average saccade rate of two monkeys. Both humans and monkeys show a similar saccadic rate in each condition, but monkeys exhibit a second noticeable peak after the target disappearance in simultaneous presentation.

**Supplementary Figure 4.**
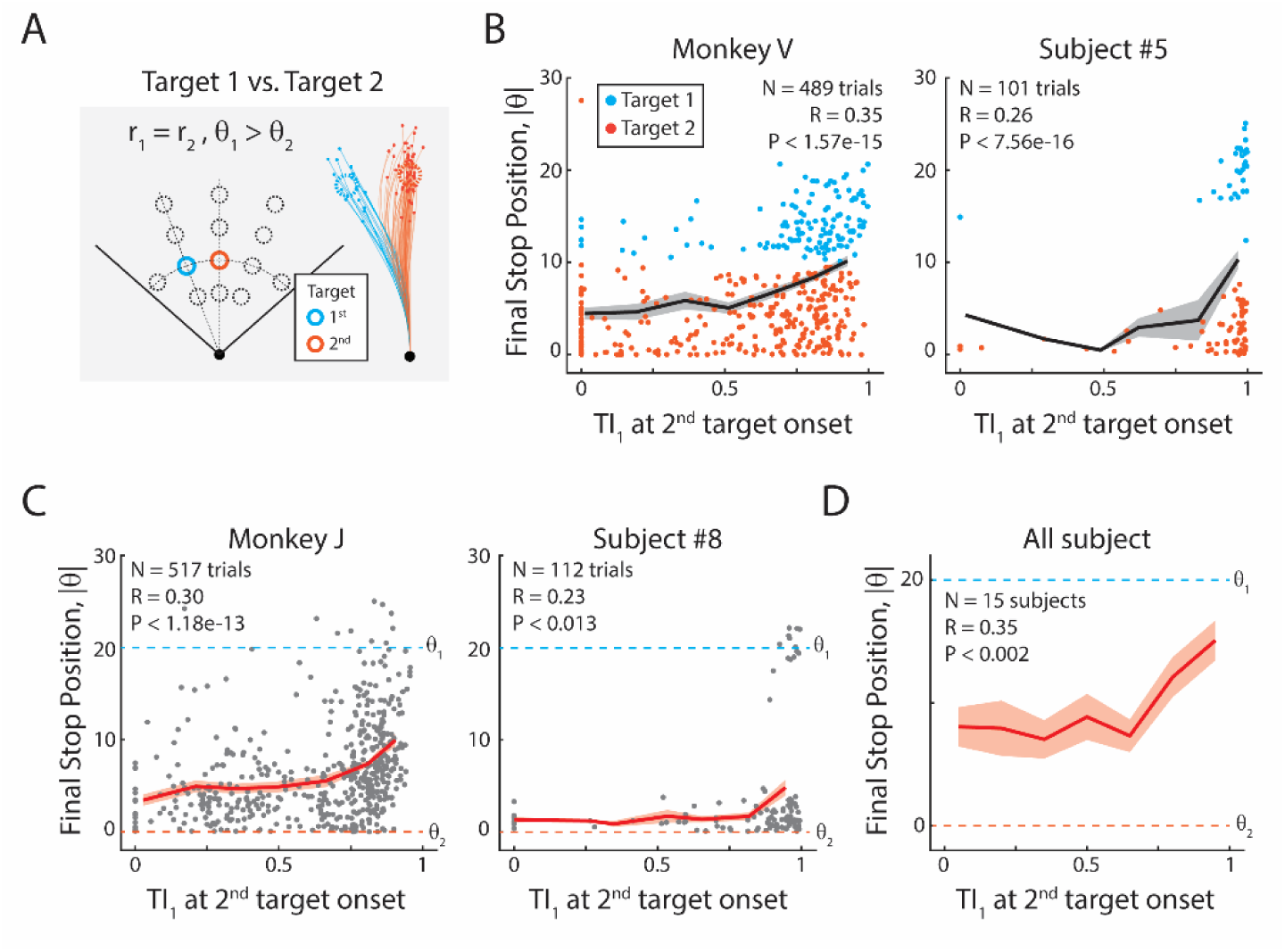
Influence of Target Tracking on Target Selection and Stop Position, related to Figure 3. (**A**) Trial conditions with sequential presentations where two targets have the same distance (r_1_ = r_2_) but differ in angle (θ_1_ < θ_2_). Sample trajectories are colored based on the final selected target. (**B**) Correlation between TI_1_ at second target onset and final stop position, |θ|, of monkey V (left) and one subject (right), colored based on the selected target. The black line denotes the average curve. (**C**) Same as B, but of monkey J (left) and another sample subject (right), without dissociating the selected target. The dashed cyan and orange lines indicate the angles of targets 1 and 2, respectively. Note that trials with high TI_1_ is correlated with closer final stop position to target θ_1_. (**D**) Average TI_1_ vs. |θ| across 15 subjects, demonstrating a significant positive correlation (Pearson’s correlation, R = 0.35, P < 0.002, N = 15).

**Supplementary Figure 5.**
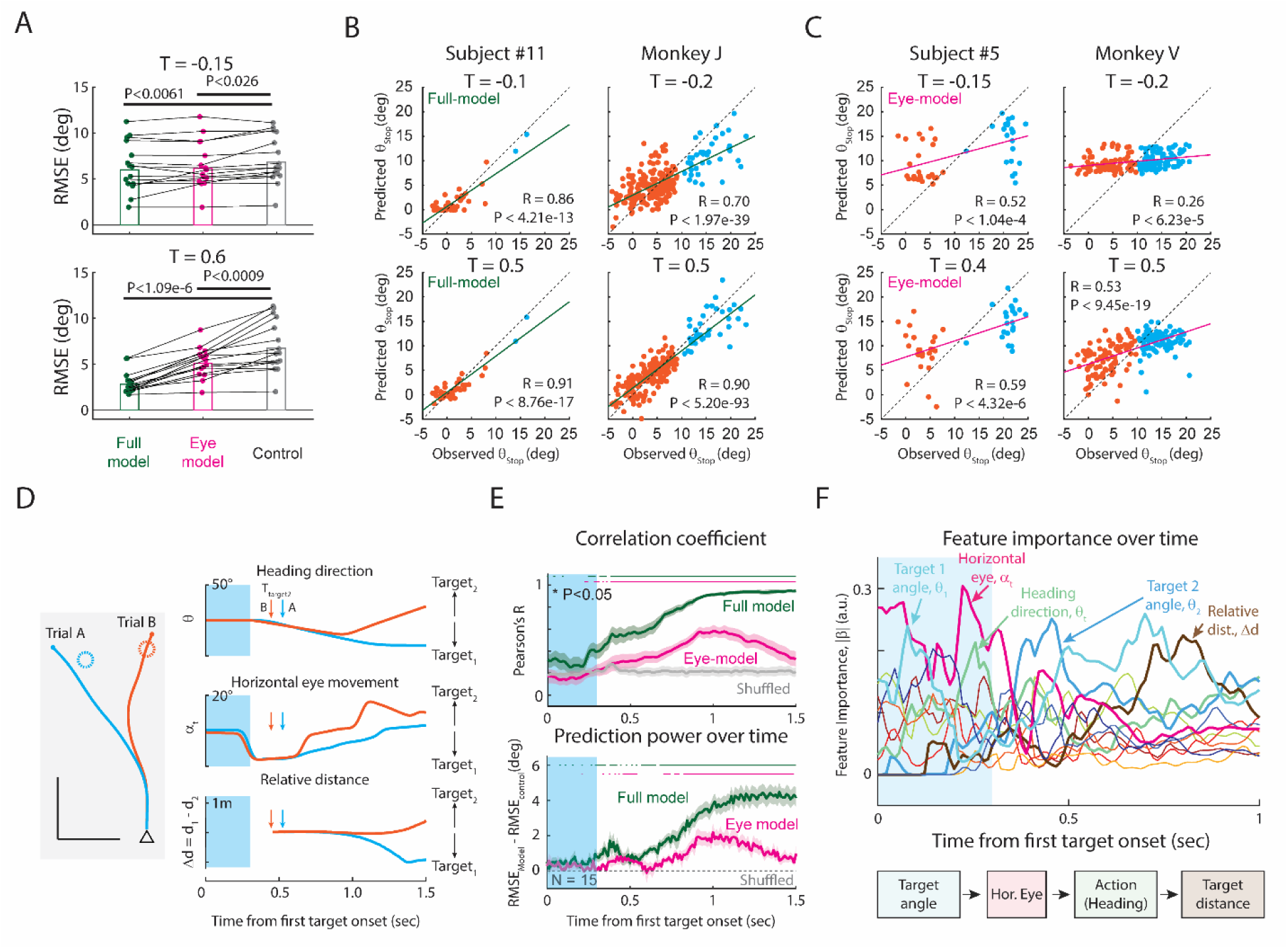
Regression Model Details, related to Figure 5. **(A)** Root-mean-square-error (RMSE) for the full regression model (green), eye-only model (magenta), and control model at T = 150 ms before (top) and at T = 600 ms after (bottom) the second target’s presentation. Both the full and eye-only models significantly outperform the control model at both times (P < 0.0034 for full model, P < 0.016 for eye-model at T = −0.15s; P < 1.34 × 10^−5^ for full model and P < 0.0018 for eye model at T = 0.6s. Paired t-test with N = 15). **(B)** Full regression model’s predicted θ_stop_ versus observed θ_stop_ before and after the second target’s presentation, demonstrating significant predictability. Pearson’s correlation was computed between the observed and predicted θ_stop_. **(C)** The eye-only model’s predicted θ_stop_ versus observed θ_stop_. The eye-only model significantly captures the tendency to stop in the leftward or rightward direction, as shown by a significant positive correlation. **(D)** Latent variables plotted against first target onset. The timing of the second target’s onset is indicated by arrows. **(E)** Correlation coefficient (R, top) and prediction power (RMSE_model_ – RMSE_shuffled_, bottom) for both models. The eye-model shows significant prediction accuracy at two different timings: 1) ∼500 ms and 2) ∼1,000 ms after first target onset. The colored line above the figures denotes the significance of each model (P<0.05, paired t-test against the shuffled control, N = 15). **(F)** Feature importance over time, indicating initial dominance of target angle and horizontal eye position in predicting stop position, followed by heading direction and target distance’s increasing importance.

## References

1. De Lafuente, V., Jazayeri, M. & Shadlen, M. N. Representation of Accumulating Evidence for a Decision in Two Parietal Areas. J. Neurosci. 35, 4306–4318 (2015).

2. Gold, J. I. & Shadlen, M. N. The Neural Basis of Decision Making. Annu. Rev. Neurosci. 30, 535–574 (2007).

3. Hanks, T. D. et al. Distinct relationships of parietal and prefrontal cortices to evidence accumulation. Nature 520, 220–223 (2015).

4. Kim, J.-N. & Shadlen, M. N. Neural correlates of a decision in the dorsolateral prefrontal cortex of the macaque. Nat. Neurosci. 2, 176–185 (1999).

5. Mante, V., Sussillo, D., Shenoy, K. V. & Newsome, W. T. Context-dependent computation by recurrent dynamics in prefrontal cortex. Nature 503, 78–84 (2013).

6. Redish, A. D. Vicarious trial and error. Nat. Rev. Neurosci. 17, 147–159 (2016).

7. Shadlen, M. N. & Newsome, W. T. Motion perception: seeing and deciding. Proc. Natl. Acad. Sci. 93, 628–633 (1996).

8. Gottlieb, J., Hayhoe, M., Hikosaka, O. & Rangel, A. Attention, Reward, and Information Seeking. J. Neurosci. 34, 15497–15504 (2014).

9. Hayhoe, M. & Ballard, D. Eye movements in natural behavior. Trends Cogn. Sci. 9, 188–194 (2005).

10. Huk, A., Bonnen, K. & He, B. J. Beyond Trial-Based Paradigms: Continuous Behavior, Ongoing Neural Activity, and Natural Stimuli. J. Neurosci. 38, 7551–7558 (2018).

11. Cisek, P. & Kalaska, J. F. Neural Mechanisms for Interacting with a World Full of Action Choices. Annu. Rev. Neurosci. 33, 269–298 (2010).

12. Yoo, S. B. M., Hayden, B. Y. & Pearson, J. M. Continuous decisions. Philos. Trans. R. Soc. B Biol. Sci. 376, 20190664 (2021).

13. Brown, T. I. et al. Prospective representation of navigational goals in the human hippocampus. Science 352, 1323–1326 (2016).

14. de Cothi, W. J. Predictive maps in rats and humans for spatial navigation. Curr. Biol. 3676–3689 (2022) 10.1016/j.cub.2022.06.090.

15. Ellmore, T. M. & McNaughton, B. L. Human Path Integration by Optic Flow. Spat. Cogn. Comput. 4, 255–272 (2004).

16. Etienne, A. S., Maurer, R. & Séguinot, V. Path Integration in Mammals and its Interaction With Visual Landmarks. J. Exp. Biol. 199, 201–209 (1996).

17. Etienne, A. S. & Jeffery, K. J. Path integration in mammals. Hippocampus 14, 180–192 (2004).

18. Heinze, S., Narendra, A. & Cheung, A. Principles of Insect Path Integration. Curr. Biol. 28, R1043–R1058 (2018).

19. Kearns, M. J., Warren, W. H., Duchon, A. P. & Tarr, M. J. Path Integration from Optic Flow and Body Senses in a Homing Task. Perception 31, 349–374 (2002).

20. Stone, T. et al. An Anatomically Constrained Model for Path Integration in the Bee Brain. Curr. Biol. 27, 3069–3085.e11 (2017).

21. Alefantis, P. et al. Sensory Evidence Accumulation Using Optic Flow in a Naturalistic Navigation Task. J. Neurosci. 42, 5451–5462 (2022).

22. Lakshminarasimhan, K. J., Avila, E., Pitkow, X. & Angelaki, D. E. Dynamical latent state computation in the male macaque posterior parietal cortex. Nat. Commun. 14, 1832 (2023).

23. Lakshminarasimhan, K. J. et al. Tracking the Mind’s Eye: Primate Gaze Behavior during Virtual Visuomotor Navigation Reflects Belief Dynamics. Neuron 106, 662–674.e5 (2020).

24. Noel, J.-P. et al. Coding of latent variables in sensory, parietal, and frontal cortices during closed-loop virtual navigation. eLife 11, e80280 (2022).

25. Stavropoulos, A., Lakshminarasimhan, K. J. & Angelaki, D. E. Belief embodiment through eye movements facilitates memory-guided navigation. Nat. Commun. 16, 11243 (2025).

26. Chen, C., Murphey, T. D. & MacIver, M. A. Tuning movement for sensing in an uncertain world. eLife 9, e52371 (2020).

27. MacIver, M. A., Patankar, N. A. & Shirgaonkar, A. A. Energy-Information Trade-Offs between Movement and Sensing. PLoS Comput. Biol. 6, e1000769 (2010).

28. Lakshminarasimhan, K. J. et al. A Dynamic Bayesian Observer Model Reveals Origins of Bias in Visual Path Integration. Neuron 99, 194–206.e5 (2018).

29. Noel, J.-P., Lakshminarasimhan, K. J., Park, H. & Angelaki, D. E. Increased variability but intact integration during visual navigation in Autism Spectrum Disorder. Proc. Natl. Acad. Sci. 117, 11158–11166 (2020).

30. Stavropoulos, A., Lakshminarasimhan, K. J., Laurens, J., Pitkow, X. & Angelaki, D. E. Influence of sensory modality and control dynamics on human path integration. eLife 11, e63405 (2022).

31. Noel, J.-P. et al. Causal inference during closed-loop navigation: parsing of self- and object-motion. Philos. Trans. R. Soc. B Biol. Sci. 378, (2023).

32. Wittig, J. H. & Richmond, B. J. Monkeys rely on recency of stimulus repetition when solving short-term memory tasks. Learn. Mem. 21, 325–333 (2014).

33. Kiani, R. & Shadlen, M. N. Representation of Confidence Associated with a Decision by Neurons in the Parietal Cortex. Science 324, 759–764 (2009).

34. Burk, D., Ingram, J. N., Franklin, D. W., Shadlen, M. N. & Wolpert, D. M. Motor Effort Alters Changes of Mind in Sensorimotor Decision Making. PLoS ONE 9, e92681 (2014).

35. Resulaj, A., Kiani, R., Wolpert, D. M. & Shadlen, M. N. Changes of mind in decision-making. Nature 461, 263–266 (2009).

36. Stone, C., Mattingley, J. B. & Rangelov, D. On second thoughts: changes of mind in decision-making. Trends Cogn. Sci. 26, 419–431 (2022).

37. Van Den Berg, R., et al. A common mechanism underlies changes of mind about decisions and confidence. eLife 5, e12192 (2016).

38. Bonnen, K., Burge, J., Yates, J., Pillow, J. & Cormack, L. K. Continuous psychophysics: Target-tracking to measure visual sensitivity. J. Vis. 15, 14 (2015).

39. Pitkow, X. & Angelaki, D. E. Inference in the Brain: Statistics Flowing in Redundant Population Codes. Neuron 94, 943–953 (2017).

40. Pärnamets, P. et al. Biasing moral decisions by exploiting the dynamics of eye gaze. Proc. Natl. Acad. Sci. U. S. A. 112, 4170–4175 (2015).

41. Shimojo, S., Simion, C., Shimojo, E. & Scheier, C. Gaze bias both reflects and influences preference. Nat. Neurosci. 6, 1317–1322 (2003).

42. Krajbich, I., Armel, C. & Rangel, A. Visual fixations and the computation and comparison of value in simple choice. Nat. Neurosci. 13, 1292–1298 (2010).

43. Krajbich, I. & Rangel, A. Multialternative drift-diffusion model predicts the relationship between visual fixations and choice in value-based decisions. Proc. Natl. Acad. Sci. U. S. A. 108, 13852–13857 (2011).

44. Richardson, D. C. & Spivey, M. J. Representation, space and Hollywood Squares: looking at things that aren’t there anymore. Cognition 76, 269–295 (2000).

45. Orquin, J. L. & Mueller Loose, S. Attention and choice: A review on eye movements in decision making. Acta Psychol. (Amst.) 144, 190–206 (2013).

46. Hayhoe, M. M. & Rothkopf, C. A. Vision in the natural world. WIREs Cogn. Sci. 2, 158–166 (2011).

47. Matthis, J. S., Yates, J. L. & Hayhoe, M. M. Gaze and the Control of Foot Placement When Walking in Natural Terrain. Curr. Biol. 28, 1224–1233.e5 (2018).

