## Supplementary Figure for "Primate Gaze Behavior Reflects Change of Mind in a Naturalistic Navigation Task"

**This PDF file includes:**

Figs. S1 to S5

### Supplementary material

#### Supplementary Figure S1

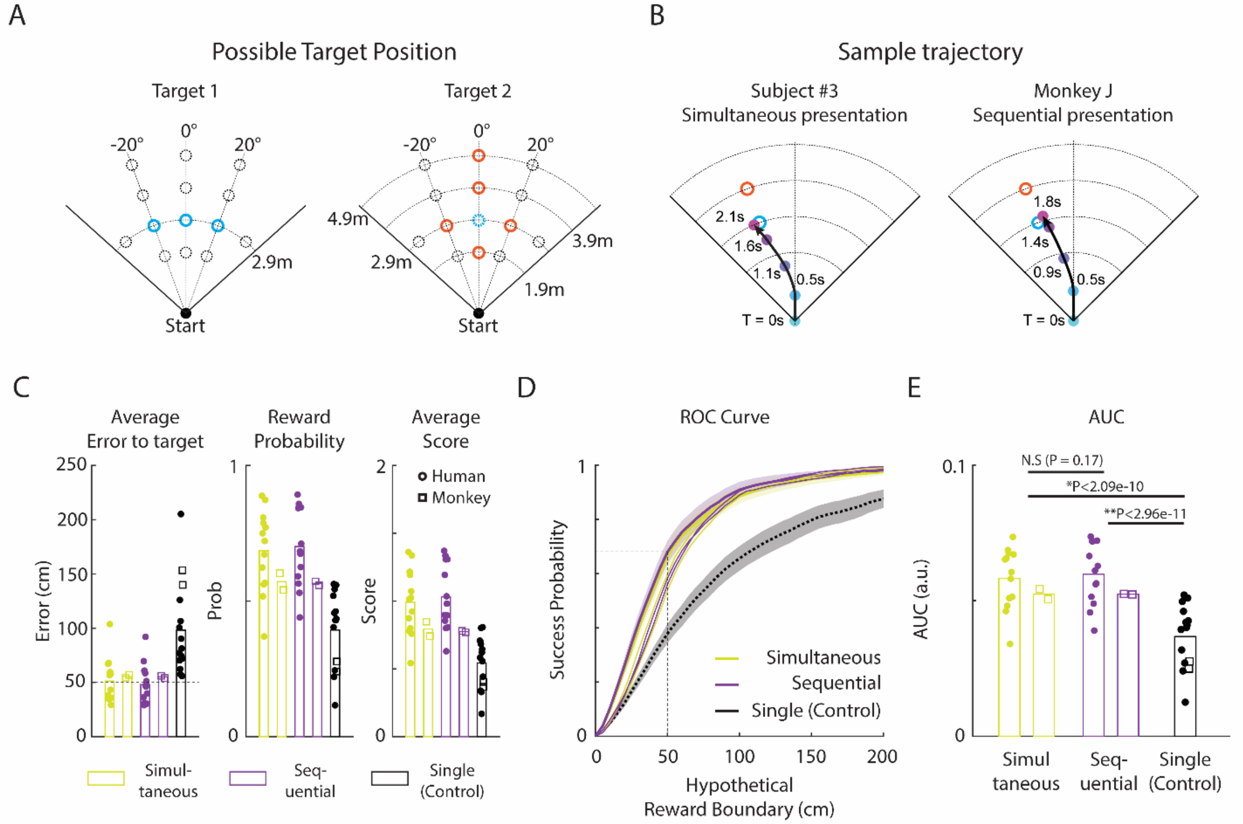


**Supplementary Figure 1. Possible Combinations of Target Presentation, Exemplary Trajectories, and Performance Comparison between Simultaneous and Sequential Presentation, related to Figure 1.**

**(A) Possible target positions**. The first target may appear at one of three positions with a 2.9 m radius at -20°, 0°, or +20° angles. The second target may appear in one of five positions: 1) the same angle but at 1.9 m, 2) 3.9 m, 3) 4.9 m, 4) the same distance but at -20°, or 5) +20° relative to the first target. **(B) Sample trajectories** of a human participant and monkey J during simultaneous and sequential presentations, respectively. Cyan and orange circles denote the first and second target positions, respectively, with black lines indicating sample trial trajectories. **(C) Performance of human and monkey subjects** under simultaneous or sequential presentation conditions. Error (left) presents the distance between the stop position vs. nearest target, averaged across all trials (rewarded and unrewarded). Dashed line shows the reward boundary (50 cm). Reward probability (middle) presents the ratio of trial with error < 50 cm. Score (right) is calculated as follows: 2 points are awarded for each trial if the error is less than 25 cm, and 1 point is given for trials where the error is between 25 cm and 50 cm. Yellow and purple dots and bars denote simultaneous and sequential presentation conditions, respectively, while black dots and bars indicate single target presentation. Filled circles represent human subjects and open rectangles represent the two monkey subjects. Humans and monkeys exhibited similar trends in all aspects, with performance in simultaneous and sequential presentations being comparable and superior to the single target condition. **(D) Receiver operating characteristic (ROC) example**, computed using a variable hypothetical reward boundary (Methods). Thick lines: Human data; ROC curve for simultaneous (yellow) and sequential (purple) conditions, with the black line representing the single target condition. Thin lines: data from two monkeys. **(E) Summary area-under-curve (AUC)** in the three conditions. Simultaneous and sequential presentations showed no significant difference in AUC performance (N.S., $P = 0.17$), but were significantly better than the single target conditions (*$P < 2.09 \times 10^{-10}$, ** $P < 2.96 \times 10^{-11}$, for simultaneous vs. single and sequential vs. single, respectively. Paired t-test, $N = 15$; 13 humans and 2 monkeys).

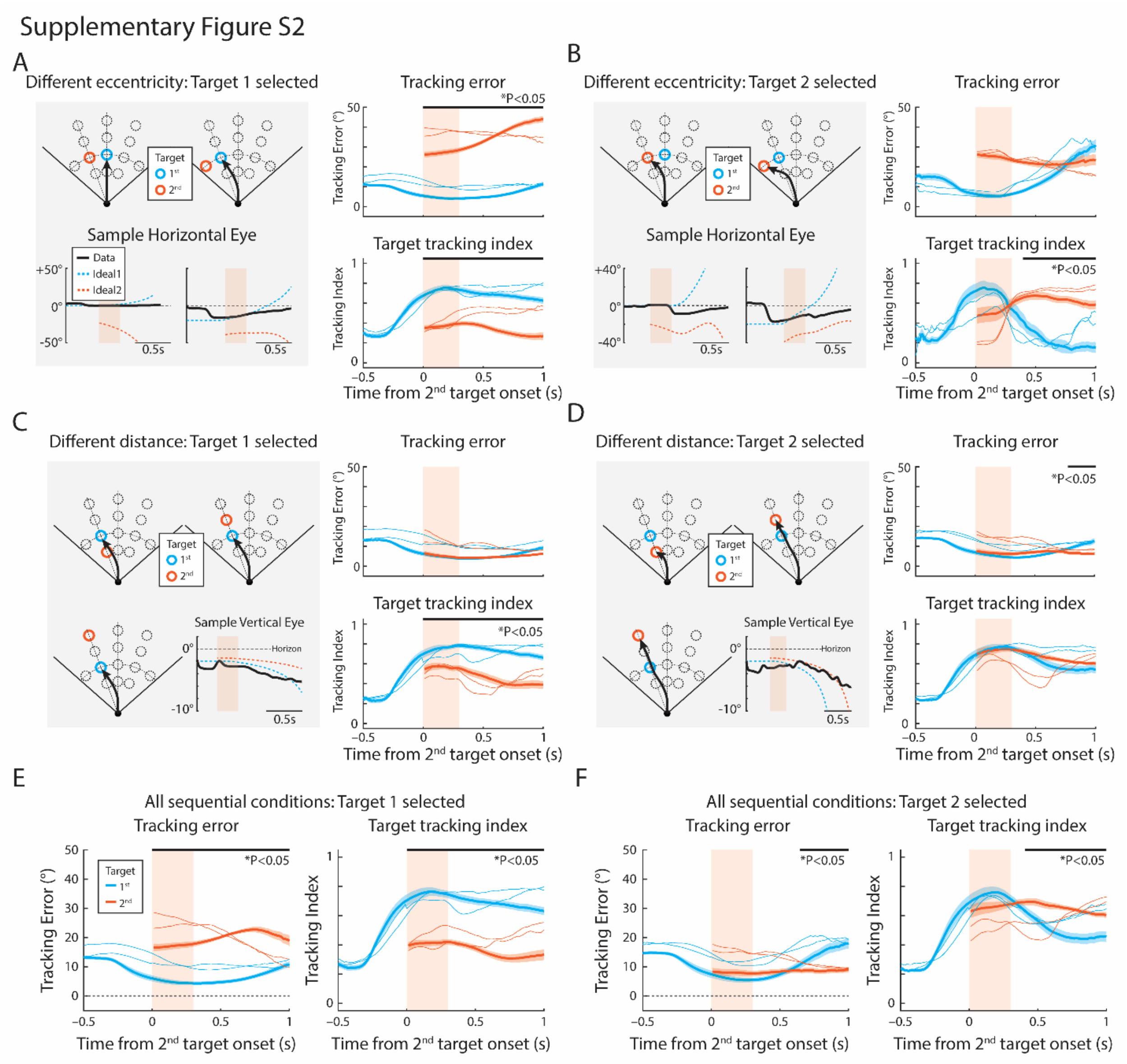


**Supplementary Figure 2. Target Tracking Error and Target Tracking Index under all Sequential Presentation Conditions, related to Figure 3.**

**(A-B)** Target tracking error and target tracking index when the second target is presented more laterally (e.g., ($\theta_1$, $\theta_2$) = (0°, ±20°) and ($\theta_1$, $\theta_2$) = (±20°, ±40°)). **(A)** When subjects select the first, less-laterally positioned target, the horizontal eye movement follows the first target position (left; cyan dotted line) rather than the second target (orange dotted line). The right panel shows that tracking error $E_1$ is significantly smaller than $E_2$ (top) and tracking index $TI_1$ is significantly higher than $TI_2$ (bottom) for all time points. **(B)** When subjects select the second, eccentric target, the horizontal eye movement shifts toward the more lateral target (bottom left; orange dotted line). Tracking error $E_2$ decreases over time (top right), and the target tracking index $TI_2$ increases significantly approximately 390 ms after the second target onset. **(C-D)** Target tracking error and target tracking index when the second target is presented at the same angle but different distance (e.g., $\theta = \theta_1 = \theta_2$, $r_2$= 1.9 m or 3.9 m or 4.9 m). **(C)** When subjects select the first target at 2.9 m, the vertical eye movement follows target 1 (bottom left; cyan dotted line) rather than target 2 (orange dotted line). The tracking error E1 and E2 were not significantly different, but the target tracking index shows $TI_1$ was significantly higher than $TI_2$ for all time points. **(D)** When subjects select the second target with a different distance, the vertical eye movement follows target 2 (bottom left; orange dotted line) rather than target 1 (cyan dotted line). Target tracking error (top right) $E_2$ becomes significantly smaller ~780 ms after target 2 presentation, and target tracking index $TI_2$ becomes subtly larger than $TI_1$ although not significant, presumably due to low variance in vertical eye movement. **(E-F)** Target tracking error and tracking indices for all sequentially presented trials, when **(E)** subject selects target 1 and **(F)** subject selects target 2. **(E)** When a subject chose target 1, $E_1$ is significantly smaller than $E_2$, and $TI_1$ is significantly higher than $TI_2$ at all time point. **(F)** When subject chose target 2, $E_2$ becomes significantly higher than $E_1$ approximately 660 ms after target 2 onset, and target tracking index $TI_2$ becomes significantly higher than $TI_1$ approximately 400 ms after target 2 onset. Shaded area denotes the standard error of human subjects, and thin lines denotes the monkey representation. All paired t-tests were conducted with N = 15.

#### Supplementary material

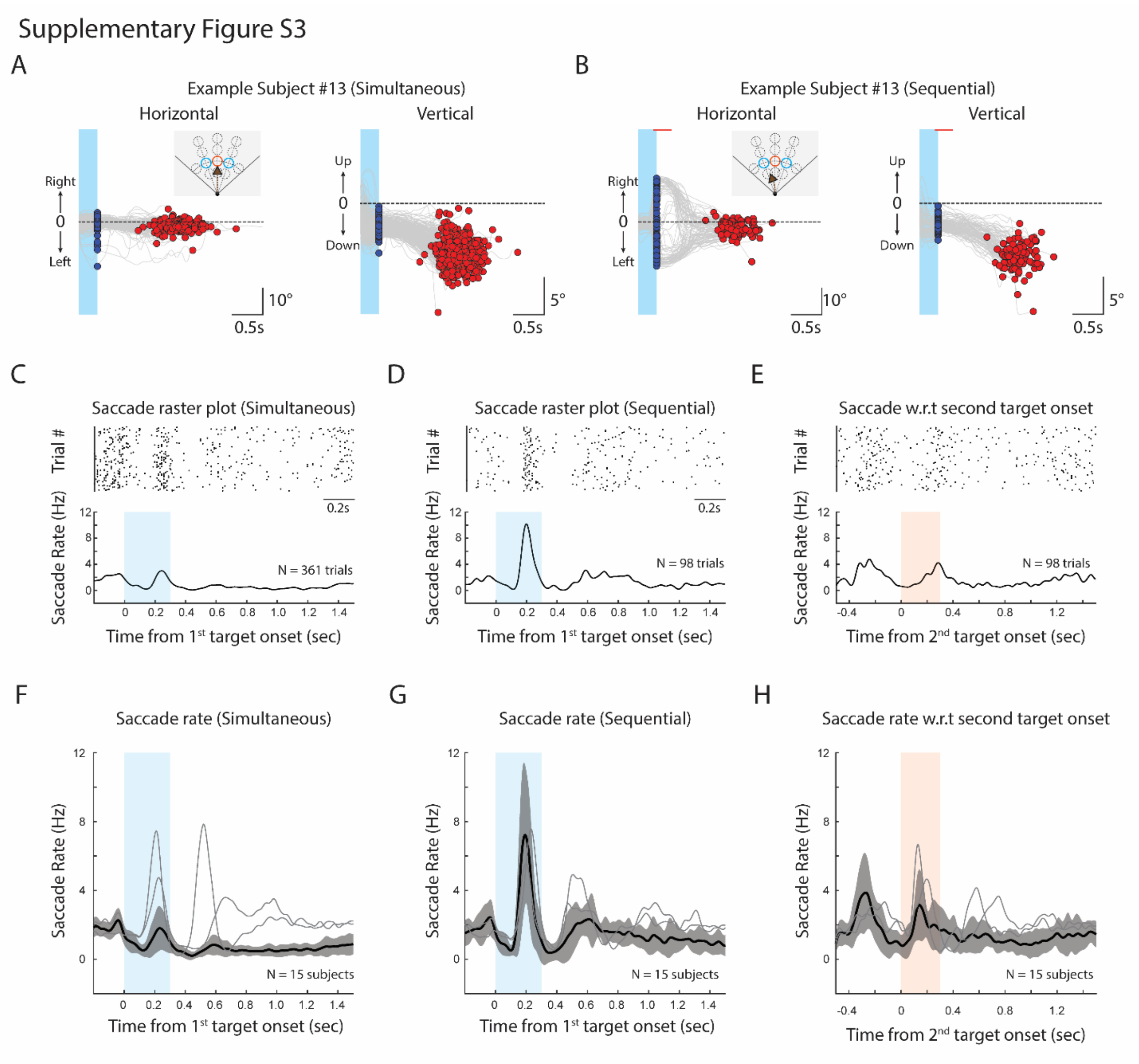


**Supplementary Figure 3. Eye movements during the task, related to Figure 3**. **(A, B)** Sample human subject's horizontal and vertical eye movements during **(A)** simultaneous and (**B**) sequential presentations when the two targets are positioned at the same distance but different eccentricity (**Figure 2F**). Blue dots denote gaze position when the target disappears, and red dots denote the final gaze position 100 ms before the subject stopped. **(A)** During simultaneous presentation, because the subject mostly chooses the central target, horizontal eye position does not make noticeable shift, but vertical eye position shifts downwards as the subject gets closer to the target. **(B)** During sequential presentation, because the target appears eccentrically, subjects make a lateral saccade, and their gaze shifts centrally as subject navigate, as shown previously [23]. **(C, D)** Sample saccade raster plot and saccade rate aligned to $1^{st}$ target onset, during **(C)** simultaneous and **(D)** sequential presentation, in trials of panel A and B. Compared to the simultaneous condition, the first eccentric target during sequential presentation induces significantly more frequent and consistent saccades. **(E)** Sample raster plot and saccade rate aligned to the second target onset during sequential presentation. **(F-H)** Average saccade rate during **(F)** simultaneous, **(G)** sequential presentation, aligned to $1^{st}$ target onset, or **(H)** aligned to $2^{nd}$ target onset, respectively. Thick line: average saccade rate of 13 human subjects; Thin lines: average saccade rate of two monkeys. Both humans and monkeys show a similar saccadic rate in each condition, but monkeys exhibit a second noticeable peak after the target disappearance in simultaneous presentation.

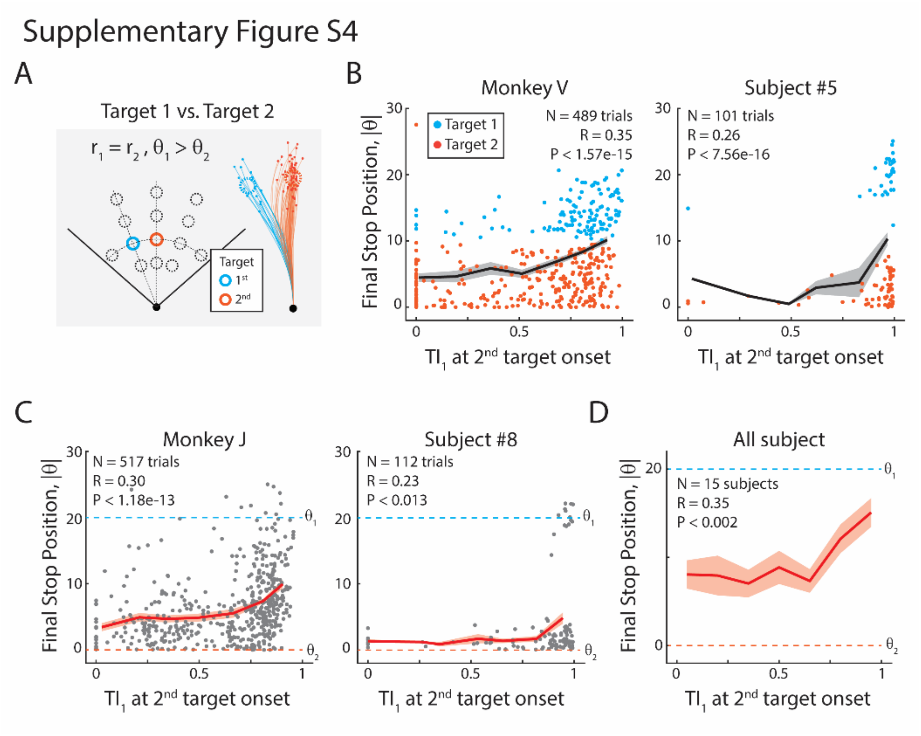


**Supplementary Figure 4. Influence of Target Tracking on Target Selection and Stop Position, related to Figure 3.**

(A) Trial conditions with sequential presentations where two targets have the same distance ($r_1 = r_2$) but differ in angle ($\theta_1 < \theta_2$). Sample trajectories are colored based on the final selected target. (**B**) Correlation between $TI_1$ at second target onset and final stop position, $|\theta|$, of monkey V (left) and one subject (right), colored based on the selected target. The black line denotes the average curve. (**C**) Same as B, but of monkey J (left) and another sample subject (right), without dissociating the selected target. The dashed cyan and orange lines indicate the angles of targets 1 and 2, respectively. Note that trials with high $TI_1$ is correlated with closer final stop position to target $\theta_1$. (**D**) Average $TI_1$ vs. $|\theta|$ across 15 subjects, demonstrating a significant positive correlation (Pearson's correlation, $R = 0.35$, $P < 0.002$, $N = 15$).

#### Supplementary material

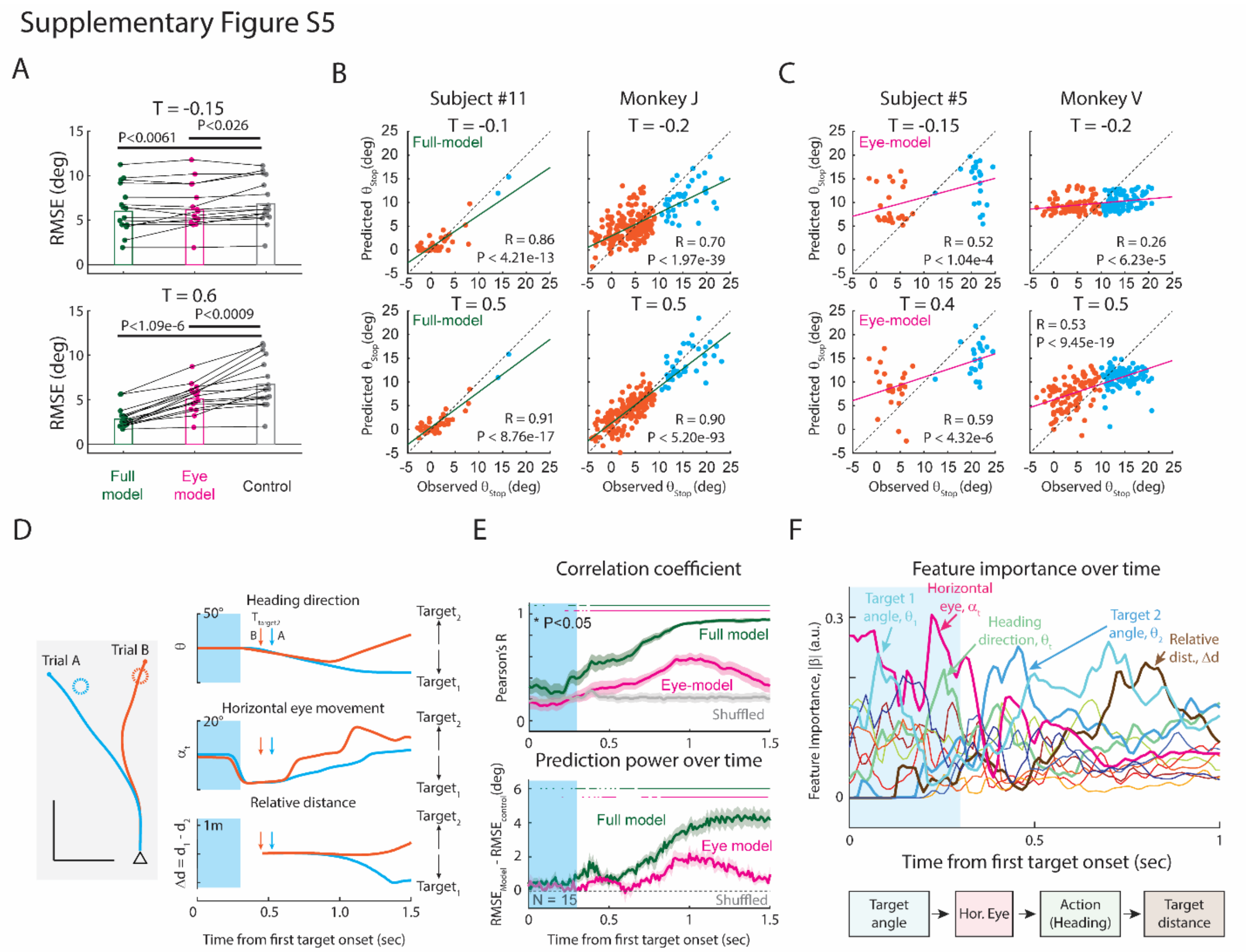


**Supplementary Figure 5. Regression Model Details, related to Figure 5.**

**(A)** Root-mean-square-error (RMSE) for the full regression model (green), eye-only model (magenta), and control model at T = 150 ms before (top) and at T = 600 ms after (bottom) the second target's presentation. Both the full and eye-only models significantly outperform the control model at both times ($P < 0.0034$ for full model, $P < 0.016$ for eye-model at T = -0.15s; $P < 1.34 \times 10^{-5}$ for full model and $P < 0.0018$ for eye model at T = 0.6s. Paired t-test with N = 15). **(B)** Full regression model's predicted $\theta_{stop}$ versus observed $\theta_{stop}$ before and after the second target's presentation, demonstrating significant predictability. Pearson's correlation was computed between the observed and predicted $\theta_{stop}$. **(C)** The eye-only model's predicted $\theta_{stop}$ versus observed $\theta_{stop}$. The eye-only model significantly captures the tendency to stop in the leftward or rightward direction, as shown by a significant positive correlation. **(D)** Latent variables plotted against first target onset. The timing of the second target's onset is indicated by arrows. **(E)** Correlation coefficient (R, top) and prediction power ($RMSE_{model}$ – $RMSE_{shuffled}$, bottom) for both models. The eye-model shows significant prediction accuracy at two different timings: 1) ~500 ms and 2) ~1,000 ms after first target onset. The colored line above the figures denotes the significance of each model ($P<0.05$, paired t-test against the shuffled control, $N = 15$). **(F)** Feature importance over time, indicating initial dominance of target angle and horizontal eye position in predicting stop position, followed by heading direction and target distance's increasing importance.
